# Inhibitory control of speech production in the human premotor frontal cortex

**DOI:** 10.1101/2023.03.01.530698

**Authors:** Lingyun Zhao, Alexander B. Silva, Garret L. Kurteff, Edward F. Chang

**Affiliations:** Department of Neurological Surgery, San Francisco, California, 94158; Weill Institute for Neurosciences, San Francisco, California, 94158; Medical Scientist Training Program, University of California, San Francisco, California, 94158; Graduate Program in Bioengineering, University of California, Berkeley, California, 94720, & University of California, San Francisco, California, 94158

## Abstract

Natural speech is full of starts and stops. Here, we studied the neural mechanisms that underlie the inhibitory control of speech, specifically the ability to stop speaking on demand. We recorded direct cortical activity while participants made continuous speech production and were given a visual cue to stop speaking. Neural recordings revealed activity in the premotor frontal cortex associated with speech stopping. Cortical sites showing stop activity were largely distinct from sites involved in active speech production or, more specifically, encoding articulatory movements. Electrocortical stimulation mapping at many premotor sites with stop activity caused involuntary speech arrest, an immediate inability to speak or vocalize. Furthermore, many speech arrest sites did not co-localize with neural activity correlating with speech motor planning or execution, contrary to this long-assumed function in clinical brain mapping. Together, these results suggest a previously unknown premotor cortical network that underlies the inhibitory control of speech, which has significant implications for understanding the dynamics of normal and altered speech production, as well as clinical brain mapping.

## Introduction

Normal conversations involve not only fluent speech production but also the ability to volitionally stop speaking quickly. An important function for the daily use of speech is the capacity to make sudden stops in the middle of an ongoing production before the intended utterance is finished (Ladefoged et al., 1973). We refer to this as “early stopping”. The control for such stopping requires suppression of planned actions and is thus believed to involve an inhibitory mechanism (Aron, 2007). Deficits in inhibitory functions have been hypothesized to cause impairments in speech and communication such as stuttering (Chang and Guenther, 2020; Eggers et al., 2010; Markett et al., 2016; Neef et al., 2018; Orpella et al., 2022) and excessive self-directed speech in attention deficit hyperactivity disorder (ADHD) (Alderson et al., 2007).

Inhibitory control has been studied extensively over the past decades. For example, previous studies have elucidated the neural basis for response inhibition, the ability to suppress a planned motor response before it is executed. The neural mechanism of response inhibition has been primarily modeled around a circuit involving the right inferior frontal cortex (rIFC), pre-supplementary motor area (pre-SMA) (Aron et al., 2014; Schaum et al., 2021; Swann et al., 2009, 2012), and several subcortical regions forming a cortico-basal ganglia loop (Aron and Poldrack, 2006; Aron et al., 2007; Chen et al., 2020). This neural circuit implements stopping through signals originating in the prefrontal cortex and is considered to function at the level of cognitive control (Aron, 2007), which is a higher level function than motor planning and execution. However, studies of response inhibition almost exclusively used paradigms where stopping occurred prior to any motor output (Logan and Cowan, 1984) and focused predominantly on hand movements. It is unclear what mechanism underlies the inhibitory control of early stopping of ongoing speech production, which may or may not rely on the same circuit as in response inhibition (Cai et al., 2012; Wagner et al., 2018; Xue et al., 2008). Fluent speech production has key differences from simple manual movements, and requires precise coordination and sequencing of multiple articulators, on the order of milliseconds, to generate an array of speech sounds (Chartier et al., 2018; Guenther, 2016; Hickok, 2012). This is in contrast to a simple movement like a button press that is considered ballistic (Logan and Cowan, 1984).

We asked how early stopping during speech production is implemented in the brain and specifically, whether the premotor cortex shows inhibitory functions to facilitate stopping. Although the prefrontal and pre-SMA cortices have been implicated in the model of response inhibition, early stopping of speech may require neural processes at a level between general cognitive functions found in the prefrontal cortex and motor control functions found in the ventral sensorimotor cortices. Therefore, the premotor cortex may play an important role. One possibility is that the premotor cortex purely supports motor planning and initiation (Wise, 1985). Neural signals from the prefrontal cortex inhibit motor planning and execution during early stopping and therefore, the premotor cortex is suppressed (Kalaska and Crammond, 1995). Another possibility is that the premotor cortex has the functionality to inhibit motor execution centers. Under this hypothesis, activation is expected in the premotor cortex for early stopping. Evidence from both animal and human studies supports the existence of inhibitory functions in the premotor cortex. (Adam et al., 2022; Buch et al., 2010; Cardellicchio et al., 2021; Duque et al., 2012; Menon et al., 2001; Mirabella et al., 2011). Clinical studies also point to the premotor cortex as an important region for stopping speech. It has long been known that electrical stimulation in the premotor and prefrontal regions, including the anterior portion of the precentral gyrus, the posterior portion of the inferior frontal gyrus (IFG), and the middle frontal gyrus (MFG), may induce speech arrest, a sudden and complete cessation of speech (Chang et al., 2017; Lu et al., 2021). Traditionally, speech arrest is interpreted as an interruption of the speech motor control signals, speech planning signals (Lu et al., 2021; Penfield and Roberts, 1959), or the “final common output for speech motor plans” (Ojemann and Mateer, 1979; Tate et al., 2014). Clinically, speech arrest is also thought to represent Broca’s area (Quiñones-Hinojosa et al., 2003). However, alternative views have been proposed recently to suggest it may be due to the inhibition of speech (Filevich et al., 2012; Silva et al., 2022).

To probe the mechanism for early stopping, we used high-density electrocorticography (ECoG) to record cortical regions across frontal, parietal, temporal, and medial areas. This methodology provided extensive spatial sampling and fine temporal resolution to track millisecond level dynamics that are essential for both speech production and stopping. Participants performed a task where they were required to immediately start and stop speaking in response to visual cues. We observed neural populations across the premotor cortex with robust activations during early stopping. We next demonstrated that this activity was unique to early stopping and did not occur during the natural passive stop at the end of a phrase. Inhibitory premotor populations did not overlap with populations important for controlling articulators during production. Finally, we found an overlap between sites showing stop activity in the task and those causing speech arrest when stimulated. Together, these results provide important evidence for a distinct, causal premotor circuit for inhibitory speech motor control.

## Results

To delineate the neural process for stopping ongoing speech production, we asked participants to perform a speech production task with early stopping guided by visual cues (referred to as the “speech stopping task”). In total, 11 participants (8 left hemisphere or LH, 3 right hemisphere or RH) were included in the analysis. All participants had high-density grid coverage over the lateral premotor cortex, with some having additional frontal, parietal, or temporal coverage. A subset of participants had coverage on the medial cortical surface (4 LH, 3 RH), providing an opportunity to investigate pre-SMA activity for early stopping, in comparison to previous studies. On each trial, a visual cue indicated when participants should start and stop their speaking, where the speech production task was to recite the days of the week at a normal pace (Fig. 1A). Participants were instructed to stop speaking immediately when the stop cue was presented. The time difference between the presentation of the stop cue and the acoustic stop of speech is referred to as the “stop reaction time” (SRT). Participants finished between 62 and 104 trials in total, across 2-3 blocks. Across all participants, early stopping was successfully achieved (Fig. 1B). We included trials with SRT longer than 0.1 s for further analysis to exclude trials where participants may have stopped at or before the stop cue by coincidence.

**Figure 1.**
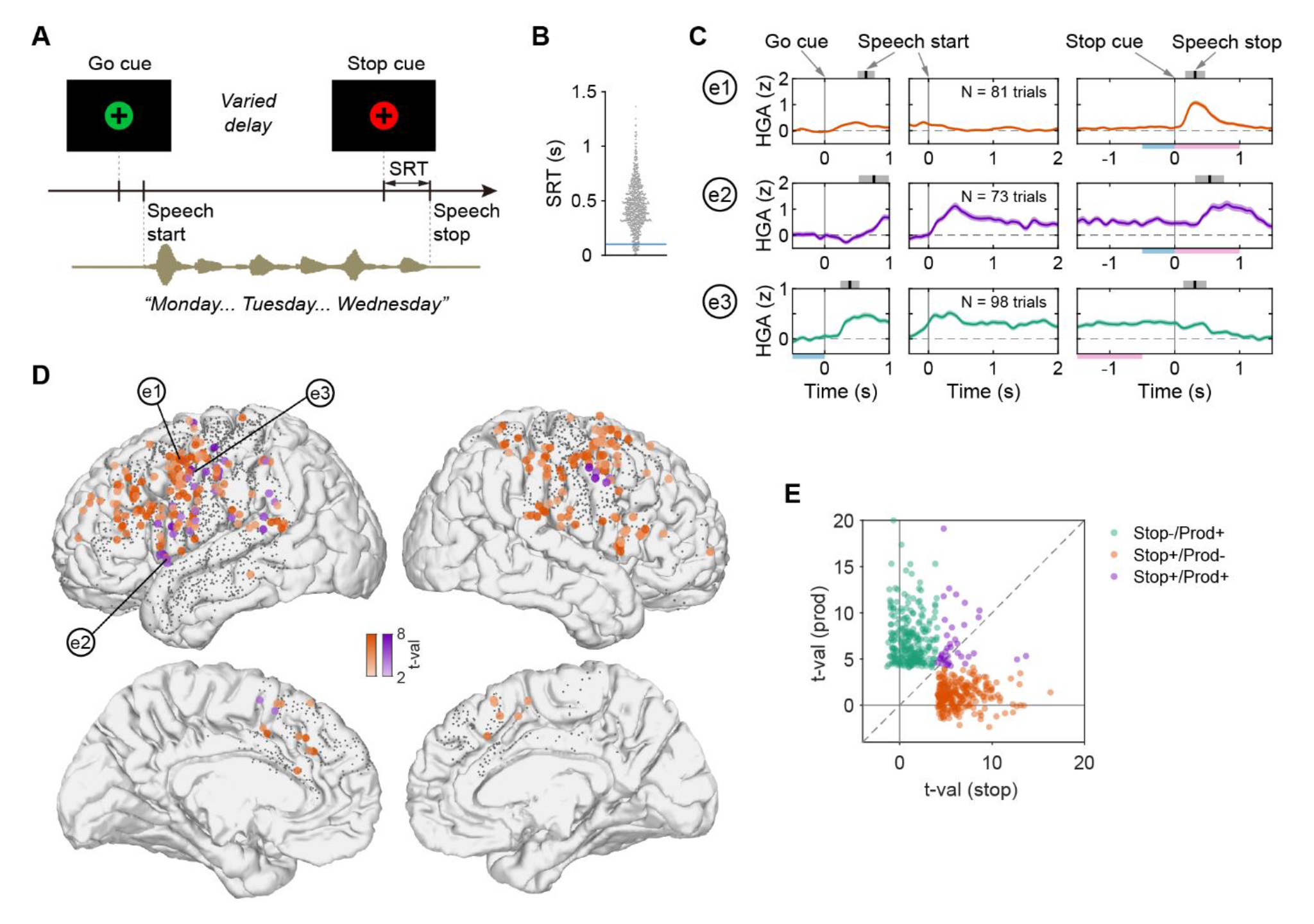
Premotor neural activation during speech stopping. (A) Schematic of the speech stopping task. The go cue is represented by a green circle. The stop cue is represented by a red circle. SRT: stop reaction time. (B) Stop reaction time (SRT) from all trials tested in all participants. Blue line: threshold for included trials (SRT ≥ 0.1s). (C) Example electrodes demonstrating three types of neural activity patterns (mean ± sem). HGA: high-gamma amplitude. Top row: electrode showing increased activity after the stop cue but no activity during production. Middle row: electrode showing increased activity during production and an additional increase after the stop cue. Bottom row: electrode showing increased activity during production but no additional activation after the stop cue. Activity is aligned to three different time points: go cue, speech start, and stop cue. The time of speech start and speech stop is marked above the panels (mean ± std). The blue bars below the panels illustrate the baseline periods. The pink bars illustrate the time periods for testing functional activity. The blue and pink bars for the top and middle rows are for testing stop activity. Those for the bottom row are for testing activity during production. (D) Location of all electrodes showing stop activity (colored circles) across participants (8 LH, 3 RH) plotted on an average brain (MNI-152). Color intensity indicates the magnitude of activity (t-value). Orange: electrodes with no production activity. Purple: electrodes with production activity. Gray dots indicate the electrode coverage. See Fig. S2A for the location of electrodes showing production activity without stop activity. (E) Scatter plot of activity magnitude during stopping and production from all electrodes showing significant activity in either condition. Each circle indicates a single electrode. Green: electrodes showing production activity but no stop activity. Orange: electrodes showing stop activity but no production activity. Purple: electrodes showing both stop activity and production activity.

### Activation of the premotor cortex for early stopping

We first asked if the premotor cortex showed activity (high-gamma amplitude [HGA]; 70-150Hz) that was time-aligned to key moments when participants were cued and executed speech production and stopping. We aligned the neural data to the go cue, speech start, and the stop cue, respectively (Fig. 1C). In an example electrode in the premotor part of the precentral gyrus, we found that HGA increased exclusively during the stopping phase of the trial (paired t-test, FDR corrected, q = 0.05, Fig. 1C top row, see Methods for details). A second example electrode in the posterior inferior frontal gyrus showed sustained activation during production (production activity) and a further increase in activity after the stop cue (Fig. 1C middle row). A third example electrode in the precentral gyrus showed increased activity during production, but no further activation after the stop cue (Fig. 1C bottom row). We refer to the increased activity after the stop cue as stop activity, regardless of whether production activity was also present. While production activity is expected in these cortical areas, the existence of stop activity suggests an inhibitory function of these cortical sites. The inhibitory function may be exclusive to a cortical site, or it can co-localize with functions serving production.

We observed a similar pattern of stop activity from each participant, with the location primarily found in the premotor frontal regions, including the ventral to middle precentral gyrus, parts of the IFG, MFG, and pre-SMA. (Fig. 1D, see also Fig. S1, S2A). To quantify the overlap between stop activity and production activity, we compared the magnitude of these activities within individual electrodes (Fig. 1E). Most electrodes showed either stop activity only (Stop+/Prod-, N = 251) or production activity only (Stop-/Prod+, N = 243). A small fraction showed both (Stop+/Prod+, N = 37, 13% of all electrodes showing stop activity). Therefore, there is little overlap between stop activity and production activity. Electrodes with stop activity but no production activity (Fig. 1D, orange circles) were located generally more anterior than electrodes with production activity but no stop activity (Fig. S2A, green circles). These results indicate there are distinct neural populations in the premotor cortex that control inhibition and motor movement of ongoing speech.

### Stop activity is specific to early stopping but not a natural finish

We asked whether stop activity reflects a volitional control of speech stops (i.e., early stopping), or whether this activity is also found when participants complete their intended speech (i.e., natural finish). A subset of participants (N = 4) completed the speech stopping task and a task in which they spoke aloud natural English sentences (Chartier et al., 2018). We first examined electrodes that were classified as Stop+/Prod- in the speech stopping task, and found examples where there was an increase in activity during early stopping, but no change in activity during the natural finish, when aligned to the time of speech stop (Fig. 2A). For all the Stop+/Prod- electrodes, the majority (80%, 43/54) were activated only in early stopping (p = 6.2×10^-8^, sign test, Fig. 2B, Fig. S3). We compared this pattern to Stop+/Prod+ electrodes and observed examples that also showed an increase in early stopping, but not in the natural finish (Fig. 2D). Similar to the Stop+/Prod- electrodes (Fig. 2A, B), the Stop+/Prod+ electrodes were more likely to show stop activity only in the early stopping case (86%, 12/14; p = 0.039, sign test, Fig. 2E, Fig. S3).

**Figure 2.**
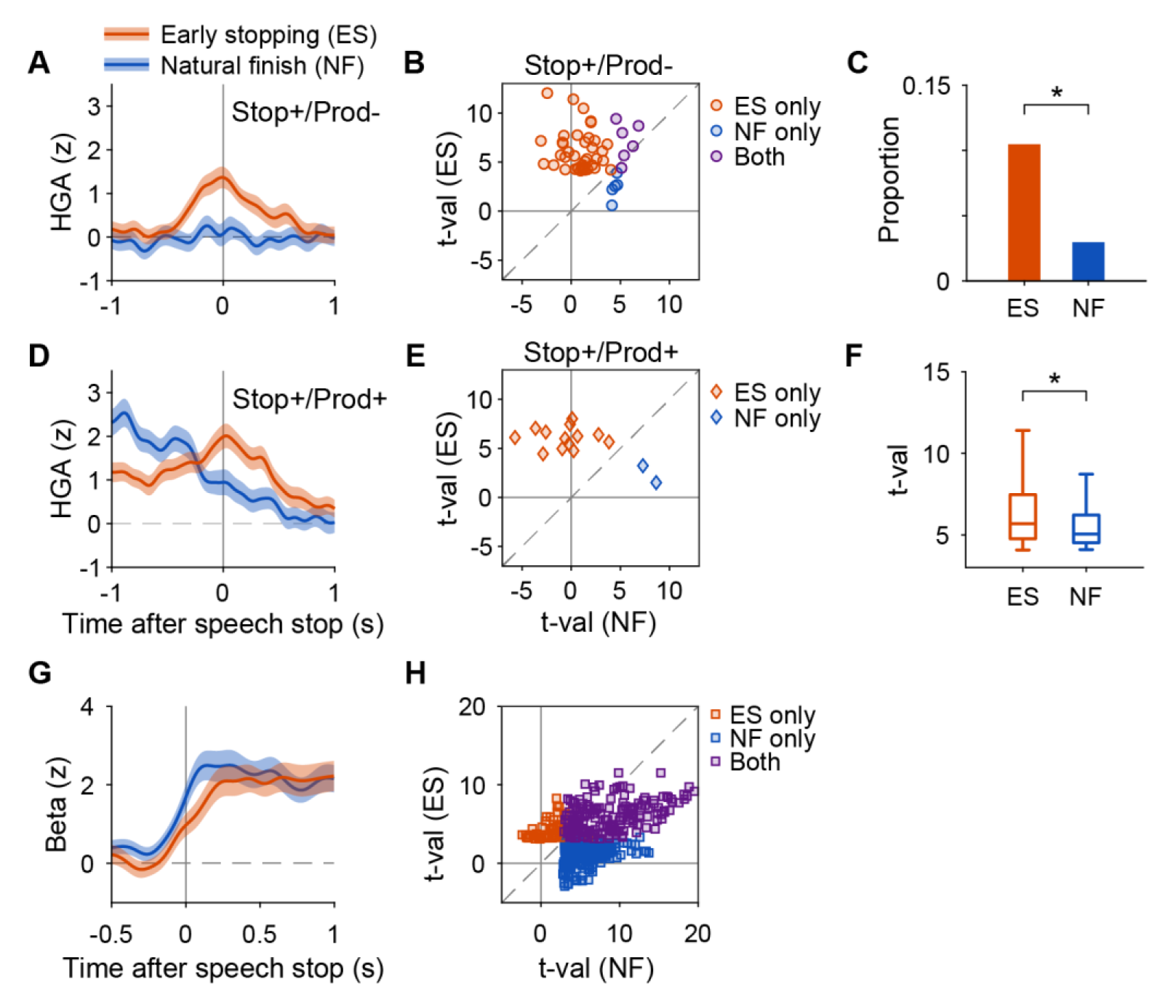
Stop activity primarily found in early stopping. (A, D) Example electrodes showing high-gamma stop activity (mean ± sem) during early stopping (ES), but not a natural finish (NF). (A) electrode with no production activity. (D) electrode with production activity. (B, E) Scatter plot of high-gamma activity magnitude during ES and NF for all electrodes showing stop activity in either condition (N = 4 participants tested both ES and NF). Each marker indicates a single electrode. (B) electrodes with no production activity. (E) electrodes with production activity. (C) Proportion of electrodes showing stop activity from 11 participants tested with ES condition and from 9 participants tested with NF condition. The asterisk indicates a significant difference. (F) Boxplot of the magnitude of stop activity in the significant electrodes from the same participants in (C). 1^st^ and 3^rd^ quartiles are marked by the borders of the boxes. Inner fences are marked by whiskers. Outliers, if any, are not plotted. The asterisk indicates a significant difference. (G) Example electrode showing increased beta-band activity (mean ± sem) during ES and NF. The location of this electrode is shown in Fig. S2C. (H) Scatter plot of beta-band activity magnitude during ES and NF for all electrodes showing increased activity in either condition (N = 4 participants tested both ES and NF). Each marker indicates a single electrode.

Since data included in this analysis were limited, we included additional participants (N = 5) with similar grid coverage who performed in the natural finish condition but not early stopping. We compared activities from all participants in early stopping (N = 11) and those from all participants in natural finish (N = 9) as two populations. We found the proportion of electrodes showing increased activity for stopping was significantly larger in the early stopping condition than in the natural finish condition (p < 1×10^-10^, chi-square test, Fig. 2C). Among these electrodes, the magnitude of activation was also larger in early stopping than in the natural finish condition (p = 0.0042, rank-sum test, Fig. 2F). These results suggest that high-gamma stop activity is predominantly found in the early stopping condition. Stop activity may be part of the volitional control process to actively inhibit ongoing speech production.

Previous studies have found beta-band activity as a neural signature for response inhibition (Swann et al., 2009). Here we tested whether the beta-band signal also showed specific activity during early stopping of speech. During motor movement, beta-band signal generally shows suppression in a wide range of sensorimotor and frontal areas. When movement finishes, beta-band activity shows an increase. In our speech stopping task, we observed increased beta-band activity around the time of speech stop (Fig. S2C). In an example electrode, beta-band activity showed an increase for both early stopping and a natural finish, after the time of speech stop (Fig. 2G). Among all electrodes showing increased beta-band activity for stopping, only a small proportion was found in early stopping but not natural finish (11%, 51/482, p = 8.9×10^-67^, sign test, Fig. 2H, orange markers), which contrast sharply to the high-gamma stop activity. This suggests that beta-band activity is not specifically related to early stopping.

### Timing of stop activity correlates with stop action

The increased activity after stop cue may be induced as a response to the cue or related to the control of stop action. To delineate the two possibilities, we performed temporal correlation analysis across single trials and found electrodes showing characteristics of these two types of functions (Fig. 3). We only included Stop+/Prod- electrodes to avoid potential confound of correlation coming from articulation signals at the end of speech. Figure 3A shows one example electrode in which the activity was found to be time-locked to speech stop (action related). To quantify this relationship, we identified the dominant activity time, an alternative measure of peak time, for the single trials that showed relatively strong activities (see Methods). Dominant activity time captures when the major event of neural activity occurs but without being biased by random fluctuations in single trials. For this electrode, there was a significant correlation between the dominant activity time and the stop reaction time (Fig. 3C). Figure 3B shows an example electrode in which the activity is found to be time-locked to the stop cue (cue related). The dominant activity time showed no correlation with the stop reaction time (Fig. 3D) and the variation of dominant activity time is small (std < 0.15 s). In total, 81 electrodes showed action related stop activity and 83 electrodes showed cue related stop activity. The remaining electrodes with stop activity were not correlated with either action or cue and were referred to as “other” (N = 87 electrodes, Fig. S4A). The location of these electrodes was similarly distributed across several brain regions, with most electrodes in the precentral gyrus and MFG (Fig. S4B, C).

**Figure 3.**
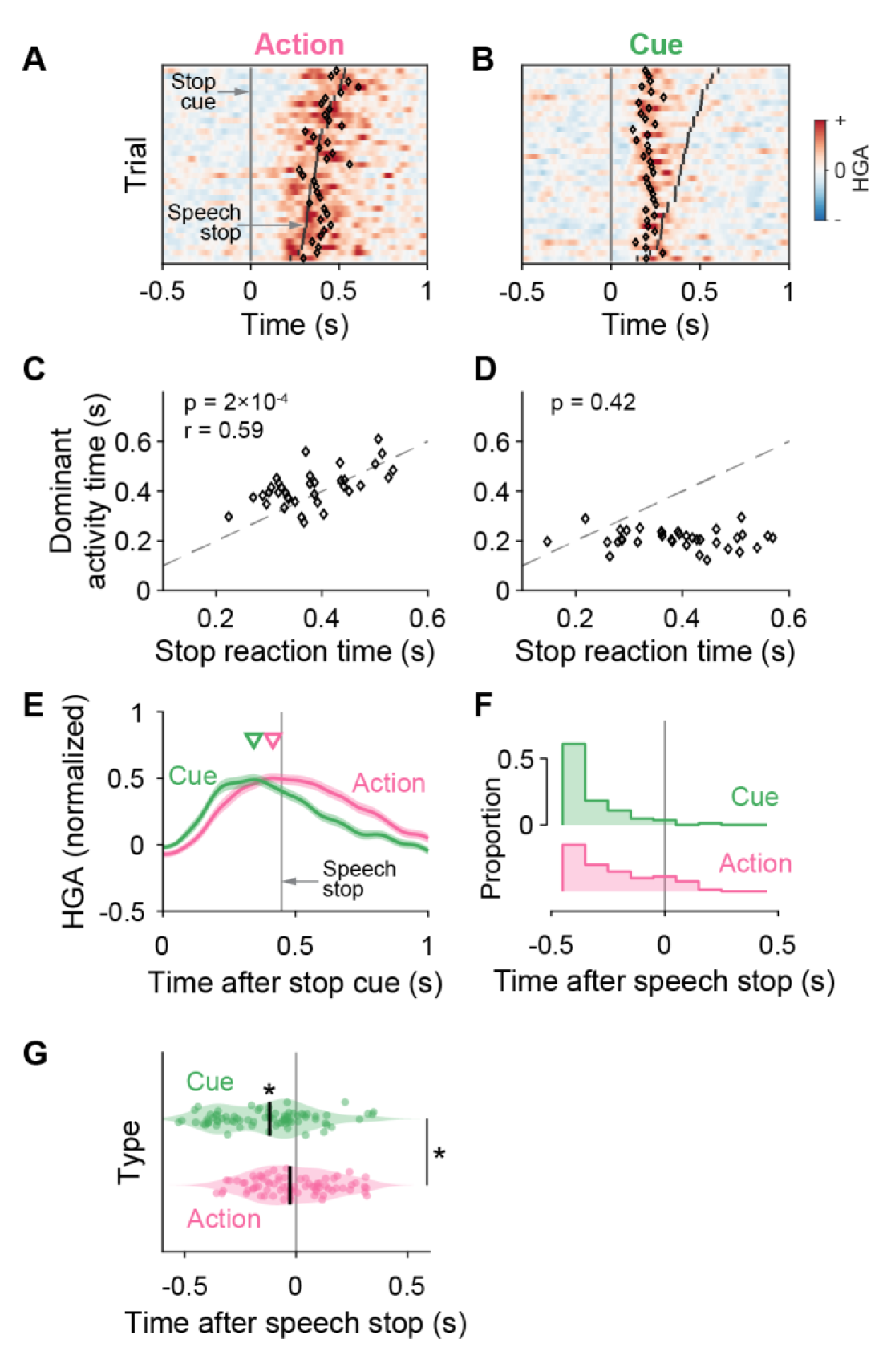
The timing of stop activity suggests a neural process leading to speech stop. (A, C) Example electrode showing activity correlated with the action of speech stopping. (A) Each row indicates a trial, sorted by the stop reaction time. The black diamond marks the dominant activity time (an alternative measure of peak time, see Methods). Only trials with strong activity where dominant activity time is identified are shown. The location of the electrode is labeled in Fig. S4B. (C) Scatter plot showing correlations between dominant activity time and stop reaction time. The dashed line is the identity line. (B, D) Similar to (A, C) Example electrode showing activity correlated to the stop cue rather than the action of speech stopping. (E) Averaged normalized activity from electrodes with cue and action related activity, time-aligned to the stop cue. Triangles: peak time in smoothed activity pattern. Vertical line: average speech stop time across participants. (F) Distribution of the activation start time for individual electrodes, relative to speech stop. (G) Distribution of the peak time for individual electrodes, relative to speech stop. Each circle indicates an electrode. Short black bars indicate medians. Asterisks indicate significant differences from time zero or between the two groups.

We next tested whether stop activity preceded and led to the stop action, which would be supported by electrodes with increases in activity before the time of speech stop but after the stop cue. We first aligned the neural activity to the stop cue and calculated the averaged normalized activity for each electrode that belongs to each type (Fig. 3E, Fig. S4D). Electrodes with both cue related and action related activity showed an increase in activity well before the time of speech stop. The cue related activity reached its peak earlier than the action related activity did. We further investigated the time of neural activity relative to the stopping action on a trial-by-trial basis. We aligned the neural activity of individual electrodes to the time of speech stop for the corresponding trial and obtained the activation start time. For the majority of electrodes, the activation start time was earlier than the time of speech stop (95% for cue related and 80% for action related activity, Fig. 3F). We also analyzed individual electrodes’ activation peak time from averaged activity aligned to the time of speech stop. The cue related electrodes showed an earlier peak time than the action related electrodes (p = 3.5×10^-4^, rank-sum test, Fig. 3G). Stop activity for most electrodes preceded the stop reaction time, suggesting this activity predicts or drives the stop action, rather than being a passive response. The difference in peak time of stop activity suggests that neural signals may track different events relevant to the stopping behavior in their temporal order. The cue related activity may be involved in identifying the need to stop speaking, which constitutes an early stage. In contrast, later stage action related activity may be involved in forming the stop command and controlling the execution of speech stopping.

### Modulation of stop activity by stopping in the middle of a word

To further understand the relationship between stop activity and the specific behavioral consequences of speech stopping, we compared instances where participants stopped speaking after the end of a word (end-of-word) and in the middle of a word (midword, Fig. 4A). A subset of participants (N = 7; 6 LH) generated sufficient midword trials and were included in this analysis. We calculated averaged high-gamma activity aligned to the time of speech stop and compared it between the midword and end-of-word stopping. We only examined the Stop+/Prod- electrodes to avoid contamination of potential articulation signal specific to stopping midword. Figure 4B shows one example electrode where the stop activity was stronger in midword trials than in the end-of-word trials (p < 0.05, cluster-based permutation rank-sum test). We found electrodes showing a similar effect across multiple participants (N = 6, all LH) and we referred to this difference in activity as stop type effect. Activity was found to be always stronger in midword trials than in end-of-word trials. The majority of electrodes (N = 13/29, Fig. 4C) with stop type effect were found in the precentral gyrus. We next asked whether this activity difference occurred before or after the speech stop. About half of the electrodes started to show activity differences before the time of speech stop (N = 15, Fig. 4D), with the precentral gyrus containing most of them (N = 10). Therefore, the additional activity found in these electrodes may signal a neural control that led to midword stopping rather than a response to the stopping behavior. At the population level, we used all Stop+/Prod- electrodes for each participant to predict whether stopping within single trials was at midword or end-of-word. A linear classifier generated predictions significantly above chance for five out of seven participants (Fig. 4E). These results indicate that stop activity distinguishes how a stop was made in each trial and further confirm that stop activity drives the behavior of speech stopping.

**Figure 4.**
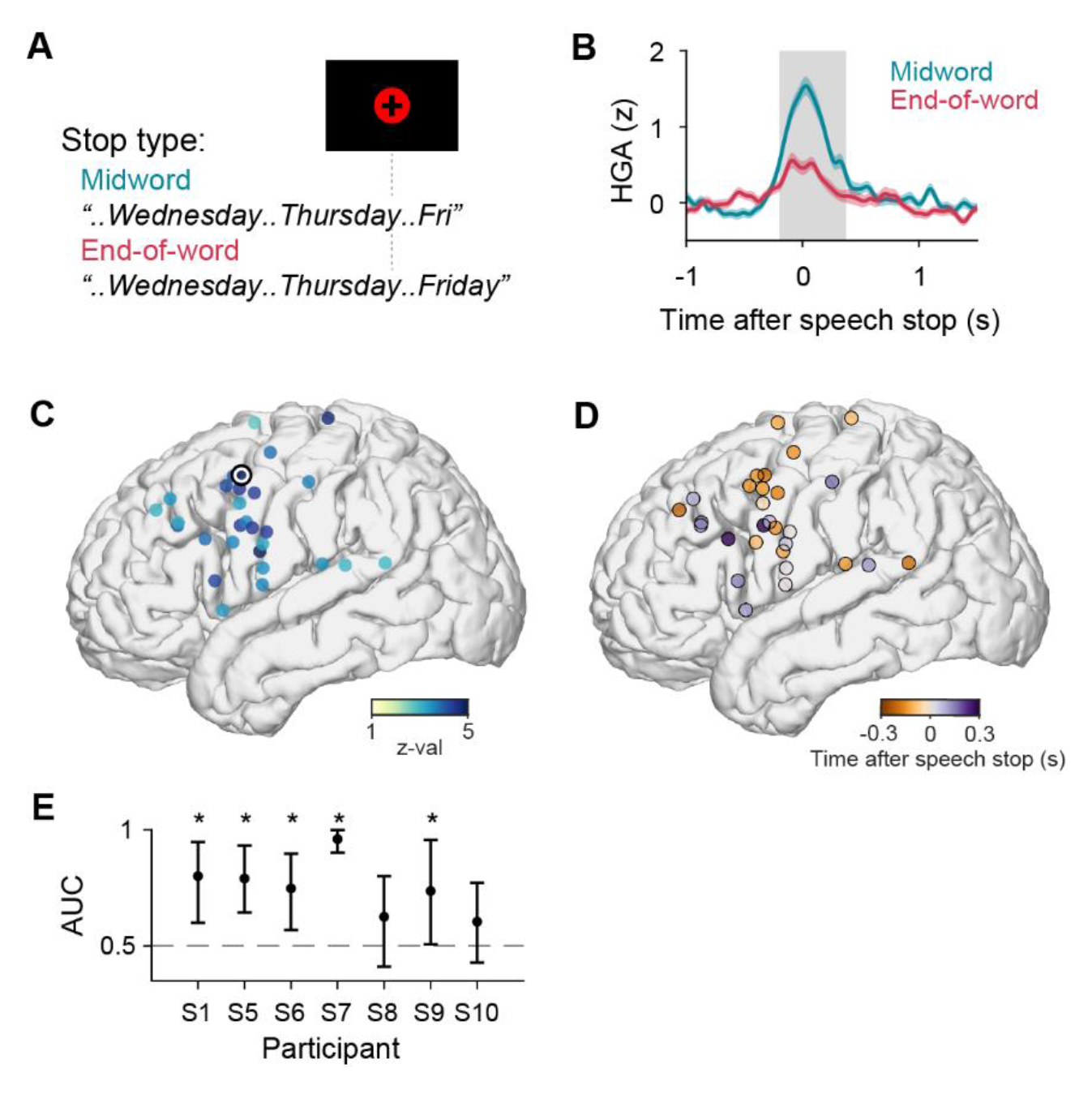
Modulation of stop activity by ending words. (A) Illustration of two types of stopping. In some trials, stopping occurred in the middle of a word (midword). In other trials, stopping occurred after finishing an entire word (end-of-word). (B) Stop activity of an example electrode (mean ± sem). Midword trials showed stronger activity than end-of-word trials did. Gray shaded region indicates the time period with significant differences. (C) Spatial location and the effect size of electrodes showing stop type effect. The black-and-white circle indicates the example electrode in (B). The one RH participant included did not show this effect. (D) Spatial location and the earliest time when electrodes started to show stop type effect, which is illustrated by the left border of the shaded region in (B). (E) Classifier performance (mean ± 95% confidence interval) for each participant where stop type was predicted by population activity of all stop electrodes. AUC: area under the ROC curve. Asterisks indicate significant differences from the chance level (dashed line).

### Relationship with other brain regions encoding speech articulation

Previous studies have found neural activity in the sensorimotor cortex that encodes the articulatory kinematic trajectory (AKT) during continuous speech production (Chartier et al., 2018). We sought to determine the relationship between stop activity and activity that controls articulation. For a subset of participants (N = 9), we were able to use acoustic-to-articulatory inversion (AAI) algorithms to infer the kinematics of vocal tract movements (see Methods). Fig. 5A shows an example electrode where its activity was strongly modulated by articulatory kinematics. We extracted 13 features to quantify the AKT and used an encoding model to fit the neural activity by these features (see Methods). Fig. 5B shows the fitted temporal filter of this example electrode, illustrating the specific articulatory kinematic pattern this electrode is sensitive to. Using held-out data, the quality of the model fit was evaluated by cross-correlation between predicted and actual activity and was found to be very high (r = 0.69). We identified electrodes encoding AKT features as those that were well fitted by AKT models (r > 0.2). Most of these electrodes were located near the central sulcus and the postcentral gyrus (Fig. 5C). In contrast, electrodes with stop activity were mostly located anterior to the AKT encoding electrodes. To further characterize the extent of overlap between the two populations, we plotted the magnitude of stop activity against the correlation coefficient in the AKT encoding models for electrodes with stop activity (Fig. 5D). Electrodes showing strong stop activity (large t-values) did not show strong encoding of AKT features (large r). This suggests that electrodes with stop activity were located in largely separate cortical regions from the AKT encoding electrodes and these two groups of activity overlapped little on the individual electrode level. This also excluded the possibility that the stop activity is simply due to the specific activation of a particular articulator during the stopping process.

**Figure 5.**
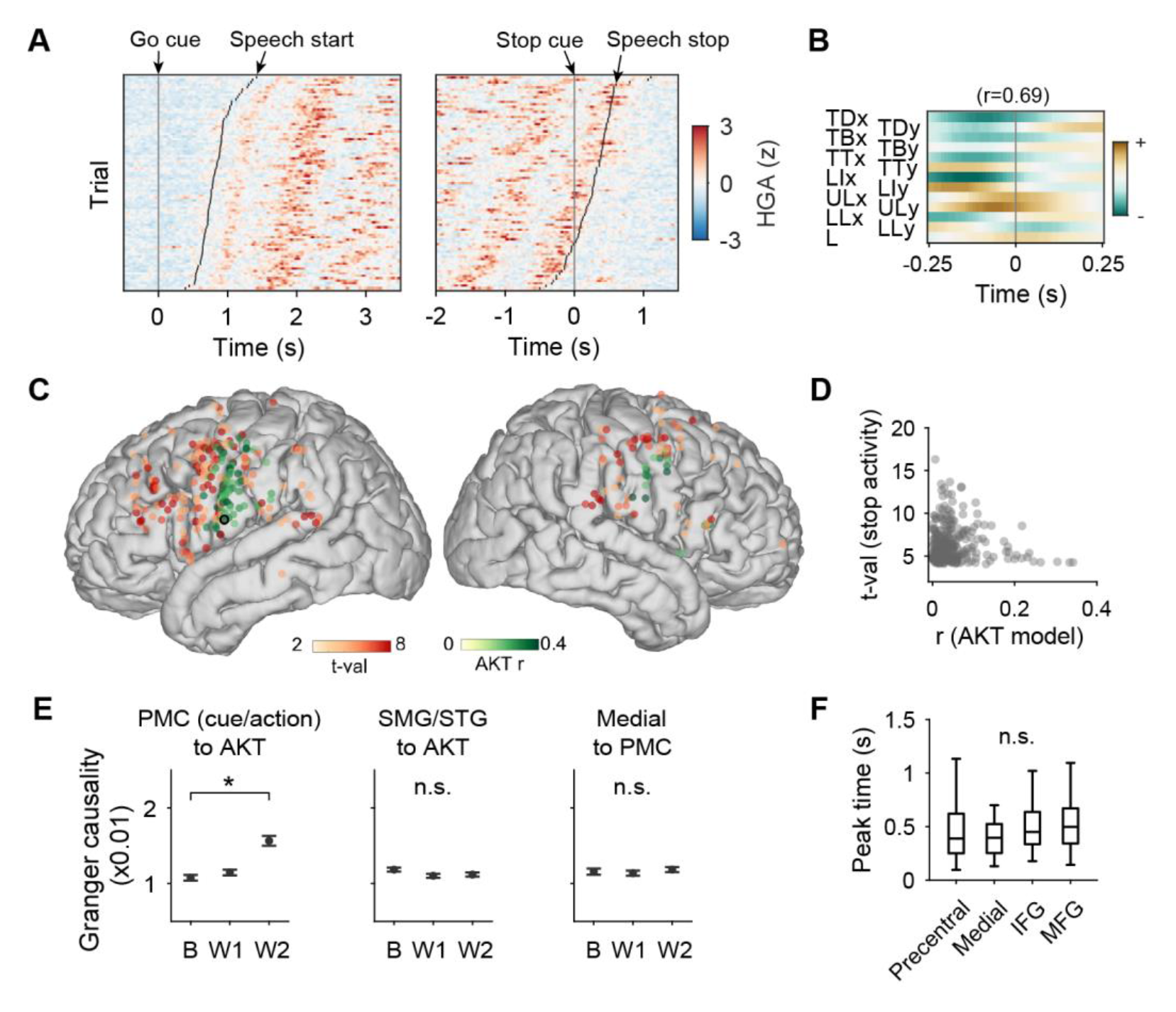
Stop activity has different spatial locations from articulatory activity. (A) Single-trial activity of an example electrode showing strong encoding of articulatory kinematic trajectory (AKT), i.e., how each articulator moves during speech. Left: trials are time-aligned to the go cue, sorted by speech start. Right: time aligned to the stop cue. The location of this electrode is indicated by a black circle in (C). (B) Temporal filter weight of this electrode for AKT features. The correlation coefficient r is indicated above the plot. TD: tongue dorsum; TB: tongue blade; TT: tongue tip; LI: lower incisor; UL: upper lip; LL: lower lip; L: larynx. (C) Spatial location of electrodes showing strong encoding of AKT features (r > 0.2, green color) and electrodes showing stop activity (red color). (D) Comparison of the magnitude of stop activity and the correlation coefficient (r) of the AKT encoding model for each electrode showing stop activity. (E) Granger causality between groups of electrodes showing stop activity and AKT encoding (mean ± sem), characterized before and after the stop cue. PMC: electrodes in the premotor cortex, including the precentral gyrus, IFG, and MFG. cue/action: electrodes with cue or action related stop activity. AKT: AKT encoding electrodes without stop activity. SMG: supramarginal gyrus. STG: superior temporal gyrus. B: baseline window [-0.5,0]s, W1: window 1 [0.0.5]s, W2: window 2 [0.5,1]s. Asterisks indicate significant differences. n.s. not significant. (F) Box plot of high gamma peak time of stop activity grouped by brain regions. 1^st^ and 3^rd^ quartiles are marked by the borders of the boxes. Inner fences are marked by whiskers. Outliers, if any, are not plotted. n.s. not significant.

Stopping in midword trials was found to have stronger stop activity than in end-of-word trials, implying that some stop activity may contain signals with specific interactions with the speech articulatory control. We next asked whether stop activity influences the AKT encoding activity for early stopping. We calculated Granger causality as a measure of directed functional connectivity before and after the stop cue based on raw neural signals between pairs of electrodes of the two groups (see Methods). We used two consecutive windows directly after the stop cue to quantify the change in Granger causality. For the electrodes in the premotor cortex with cue or action related stop activity, there was a significant increase in Granger causality towards AKT encoding electrodes in the [0.5, 1] s window (p < 0.05, repeated measures ANOVA, post hoc analysis with the Bonferroni corrections, Fig. 5E, left panel). As a control analysis, the temporal-parietal region, including the supramarginal gyrus (SMG) and the posterior superior temporal gyrus (STG), did not show a significant change in Granger causality from electrodes showing stop activity to AKT encoding electrodes (Fig. 5E, middle panel).

Lastly, we asked whether the premotor stop activity was related to or originated from neural populations in medial regions, such as pre-SMA, given prior knowledge that pre-SMA contributes to response inhibition. The Granger causality from electrodes showing stop activity in the medial regions to those in the premotor cortex did not show a significant change after the stop cue (Fig. 5E, right panel). When comparing the peak time of high-gamma activity across regions, the activity in the medial regions did not occur earlier than that in the premotor cortex. (p > 0.05, Kruskal-Wallis test, Fig. 5F). Therefore, it is unlikely that the stop activity in the premotor cortex was simply communicated from the medial cortex, such as pre-SMA. Rather, evidence points to intrinsic neural populations in the lateral premotor cortex that drive early stopping behavior through communication with articulatory populations.

### Partially overlapping location of stop activity for speech and hand movement

In the model for response inhibition, a shared neural circuit is responsible for stopping movements of different modalities (Xue et al., 2008). Here we tested whether early stopping of ongoing speech production and hand movement shared the same mechanism. A subset of participants performed a hand movement stopping task in which they pressed a button repetitively after the go cue and released it immediately after the stop cue (N = 8 participants, 7 LH, 1 RH, only LH data are included, 54-71 trials finished by each participant). We followed the same criterion to identify stop activity as for speech and used the selectivity index (SI) to quantify whether single electrodes had stop activity for speech production, hand movement, or both. To compare the spatial location, we excluded the electrodes with sustained activity during speech production or hand movement, to avoid potential bias induced by the motor homunculus. The electrodes showing stop activity for speech but not hand (“speech only”) were found towards the ventral and middle part of the precentral gyrus, IFG, and MFG regions (Fig. 6A, left panel, red circles). It is worth noting that electrodes with stop activity for both speech and hand (“both”) were also found in the ventral and middle precentral regions (Fig. 6A, left panel, blue circles). The electrodes with stop activity for hand but not speech (“hand only”) were primarily found in the dorsal part of the precentral gyrus (Fig. 6A, left panel, green circles). On the medial side, the distribution of these three types of activity did not show any spatial pattern (Fig. 6A, right panel). The distributions of “speech only” and “both” electrodes along the dorsal-ventral axis overlapped, whereas the distribution of “hand only” electrodes was largely found more dorsally (Fig. 6B). Electrodes with cue related activity constitute the highest proportion for “both” (65%). Electrodes with action related activity and the remainder of the electrodes had a proportion of “both” of 55% and 29%, respectively. This result suggests that the ventral and middle premotor areas may show an overlap of stop activity across multiple motor modalities, such as speech and hand movement, although many individual sites within these areas still show stop activity specific to speech.

**Figure 6.**
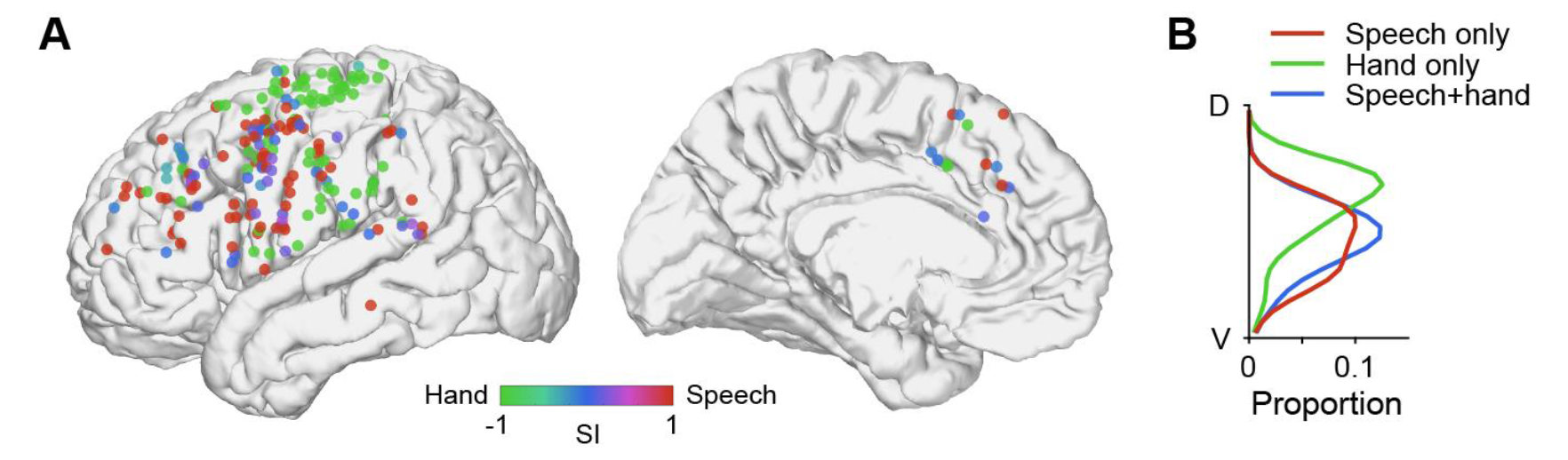
Comparison of stop activity for speech production and hand movement. (A) Location of electrodes showing stop activity to “speech only” (red), “hand only” (green), and “both” (blue). SI: selectivity index (see Methods). Only LH data are included. Electrodes with activity during production or hand movements are excluded. (B) Distribution of the location on the average brain along the dorsal-ventral axis for the three types of electrodes.

### Overlap between stop activity and stimulation induced speech arrest

In clinical practice, neurosurgical patients sometimes undergo functional brain mapping using electrocortical stimulation. Speech arrest, where a complete cessation of speech occurred during stimulation, has been traditionally interpreted as an interruption of neural signaling for speech articulation or planning (Lu et al., 2021; Penfield and Roberts, 1959). For a long time, the identification of speech arrest sites is assumed to be specific to “Broca’s Area” by many researchers and clinicians (Brannen et al., 2001; Fitzgerald et al., 1997; Ojemann, 1983; Quiñones-Hinojosa et al., 2003). However, an alternative explanation for speech arrest is that stimulation evokes an inhibitory processing mechanism that stops speech. Here, we investigate these possibilities by comparing the spatial location of stop activity with speech arrest. A subset of left-hemisphere participants with the speech stopping task underwent electrocortical stimulation through the same ECoG grid for clinical purposes (N = 7 participants). Participants counted aloud continuously while stimulation was delivered at a time unpredictable to the participants. We characterize two types of stimulation effect: speech arrest and speech error/orofacial effect (Fig. 7A). Speech arrest was identified when speech phonation was completely absent and there was no major orofacial movement. Speech error was identified when speech was able to continue during stimulation but was dysarthric, and/or associated with involuntary movements of the face or throat. At some sites, counting was not tested but passive stimulation without speech production induced motor movement or sensation around the orofacial regions. These sites are included as the orofacial effect.

**Figure 7.**
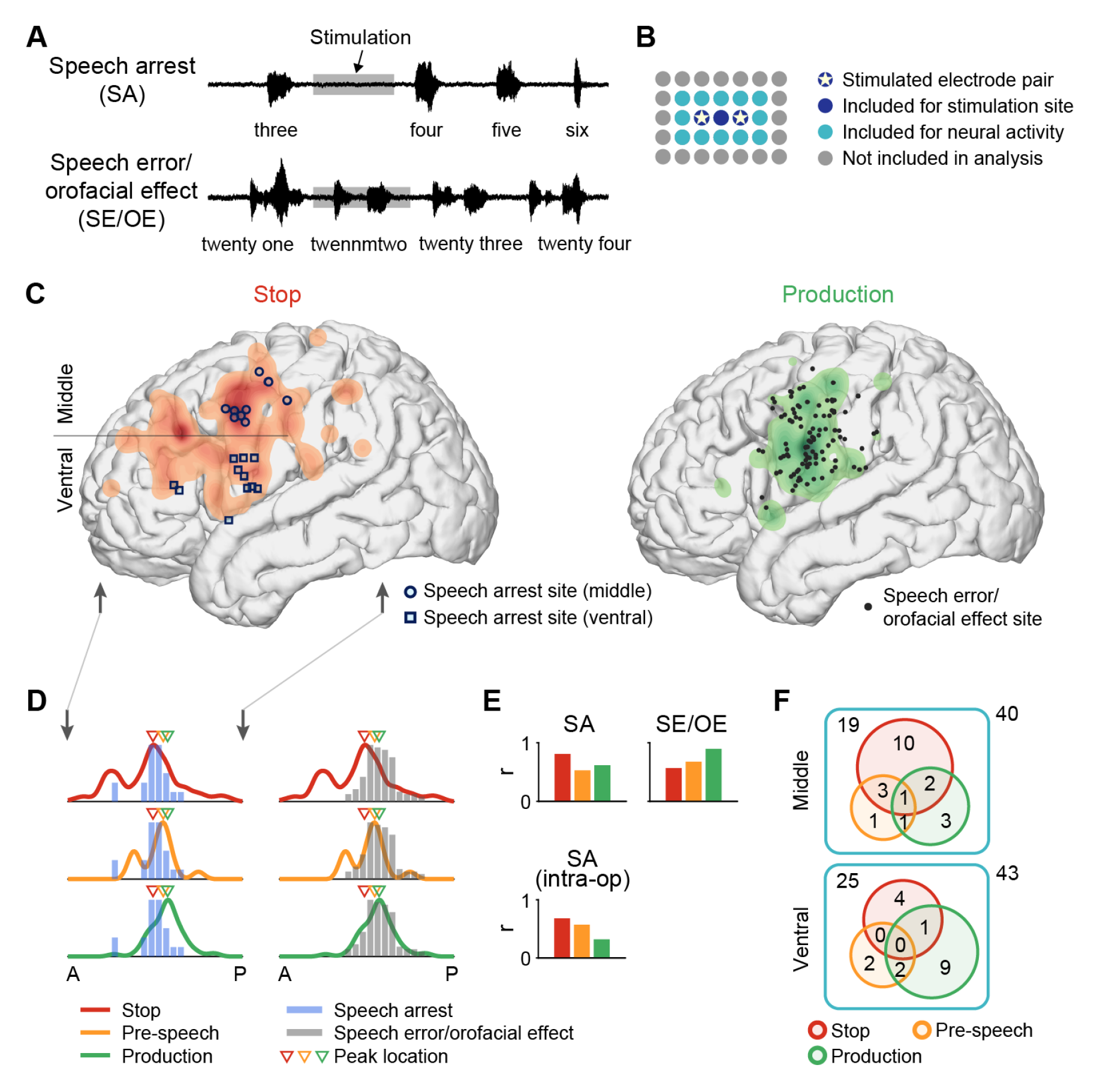
Location of speech arrest sites correlates with that of the stop activity. (A) Example waveforms of two stimulation effects on speech. The gray shaded region indicates the stimulation duration. (B) Schematic of the high-density grid. Pentagrams indicate a pair of electrodes at which bipolar stimulation was delivered. Pairs are typically separated by an additional electrode in the middle due to the limitations of the clinical stimulator. Dark blue: electrodes included as stimulation sites, used in (C, D, E). Light blue: additional electrodes within the putative current spread area. Electrodes in dark blue or light blue are included for analyzing the neural activity in (F). (C) Left: density map of electrodes with stop activity but no pre-speech or production activity, shown on an average brain, overlaid with speech arrest sites. The horizontal line separates the middle and ventral clusters of speech arrest sites. Right: density map of electrodes with production activity but no stop or pre-speech activity, overlaid with sites with speech error/orofacial effect. Vertical arrows match those in (D) and indicate the boundary of the anterior-posterior axis in (D). Activity in the temporal lobe is excluded for clarity. (D) Spatial distribution of stimulation sites (bars) and electrodes with activities (curves) in (C) along the anterior-posterior axis. Colored triangles indicate the peak locations of the spatial distribution for the three types of activities. Speech arrest sites are most likely to be found where stop activity has the highest density. Speech error and orofacial effect sites are most likely to be found where production activity has the highest density. A: anterior; P: posterior. (E) Top row: correlation coefficient between the spatial distributions of stimulation sites and electrode activities in (D). Bottom row: similar to the top row, using speech arrest sites from the intra-operative mapping dataset. The location of speech arrest sites has the best correlation with that of the stop activity. (F) Venn diagram showing overlap of stimulation sites and different types of neural activity at the individual electrode level, using electrodes at and in the vicinity of speech arrest sites. The numbers within the circles indicate electrodes showing one or more types of neural activity. The number at the upper left corner of the cyan box indicates electrodes not showing any of these activities. The number outside of the box indicates the total number of electrodes. In the middle cluster, speech arrest sites are predominantly associated with stop activity. In the ventral cluster, speech arrest sites are most often associated with production activity.

Due to the clinical stimulation setup, bipolar stimulation was usually delivered through a pair of non-adjacent electrodes in the high-density grid (Fig. 7B, pentagrams). We consider these two electrodes together with the electrode in between as a stimulation site (Fig. 7B, dark blue circles). We compared the location of stimulation sites (speech arrest: N = 7 sites, 21 electrodes from 5 participants; speech error/orofacial effect: N = 38 sites, 114 electrodes from 7 participants) to density maps of neural activity found in the speech stopping task (Fig. 7C). We did not include stimulation sites from the medial brain areas due to limited amount of data. To test our hypotheses, we consider stop activity as serving the function of inhibition, production activity as serving the function of motor articulation, and activity prior to speech onset (“pre-speech” activity) as serving the function of speech motor planning. For production activity, we restricted to sustained activity because for stimulation to interrupt motor activity during speech, one would expect there is ongoing activity to interrupt with. We compared each of the two stimulation effects to each of the three neural functions. To illustrate the results, we first examined the overlap of speech arrest sites with electrodes showing stop activity but without production or pre-speech activity. Speech arrest sites were localized within the region where stop activity was found, with a majority in the precentral gyrus (Fig. 7C, left panel). This supports the hypothesis that speech arrest may be due to inhibitory functions. The location of speech arrest sites did not correlate well with pre-speech activity (Fig. S5A). Furthermore, we found that the speech error/orofacial effect sites co-localize with electrodes showing production activity with no stop or pre-speech activity (Fig. 7C, right panel). This suggests that errors with continued production during stimulation may largely be due to motoric effects. To quantify all comparisons, we projected electrode coordinates onto the anterior-posterior axis and summarized the distribution (Fig. 7D). The peak of the stop activity distribution is found to be the most anterior among the three, which aligns best with the densest location of speech arrest sites (Fig. 7D, left column). The overall distribution of the stop activity also matches that of the speech arrest sites (Fig. 7D, top left panel). The peak of production activity distribution is found to be the most posterior among the three, which aligns the best with the densest location of speech error/orofacial effect sites (Fig. 7D, right column). The overall distribution of production activity matches that of the speech error/orofacial effect sites (Fig. 7D, bottom right panel). The peak of pre-speech activity resides in the middle and the distribution of pre-speech activity does not match well with either type of stimulation sites (Fig. 7D, middle row). The correlation between the two types of distributions further confirms this relationship. The distribution of stop activity is best correlated with that of speech arrest sites, whereas the distribution of production activity is best correlated with that of speech error/orofacial effect sites (Fig. 7E, top row). To test whether our finding based on stimulation through ECoG grids is generalizable, we studied a separate dataset from intra-operative mapping (N = 34 participants) and found a similar result. The speech arrest sites (N = 36 sites from 20 participants) found in the intra-operative mapping are in general located within the area where stop activity was found (Fig. S5B). The spatial distribution of the speech arrest sites is best correlated with that of stop activity (Fig. 7E, bottom panel).

Lastly, we compared the overlap between speech arrest sites and neural activity at the individual electrode level, to provide fine-grained information on the underlying mechanism. Previous studies have found that bipolar stimulation may generate a current spread across a distance that overlaps with adjacent electrodes in the high-density grid (Muller et al., 2018). Therefore, we included electrodes directly surrounding the stimulation site (Fig. 7B, cyan circles) in additional to those directly stimulated (Fig. 7B, dark blue circles). For an example stimulation site illustrated in Fig. 7B, a total of 15 electrodes are included in the analysis. We asked what type of neural activity these electrodes showed and grouped the speech arrest sites into two clusters according to a recent study (Lu et al., 2021). One cluster is around the ventral precentral gyrus and the other one is around the middle precentral gyrus (Fig. 7C, left panel). In total, 83 electrodes are included (43 ventral, 40 middle). In the middle cluster, electrodes at and around the speech arrest sites primarily showed stop activity, whereas, in the ventral cluster, electrodes at and around the speech arrest sites primarily showed production activity (Fig. 7F). Overall, these results support that speech arrest found in clinical stimulation mapping can be caused by inhibition, rather than an interruption of motor execution or planning of speech. Stimulation at regions with stop activity may induce an inhibitory behavioral effect, providing evidence for causal functions of the stop activity. The middle precentral gyrus is likely to be an important area for this inhibition (Silva et al., 2022).

## Discussion

In this study, we investigated the neural basis for inhibitory control of speech. Specifically, we found neural signals that implement sudden and volitional stops during ongoing speech production, which is an important function for normal and fluent verbal communication. Through high-resolution neural recordings, we identified activity in the premotor cortex that was increased after the stop cue rather than being suppressed, suggesting inhibitory functions of the premotor cortex. By comparing across task conditions and trial-by-trial variations, we found that the premotor stop activity was specific to active, volitional stopping and was absent during the natural completion of sentences. Furthermore, stop activity reflected specific components of speech stopping, including the initial response to the stop cue and the execution of the stop action itself. The magnitude of activity was modulated by whether stopping occurred in the middle of a word, suggesting that the inhibitory control may contain specific signals to interact with speech articulation. Although many stop electrodes localized to areas traditionally considered part of the speech production network, the spatial location of these electrodes was largely distinct and anterior to those encoding articulatory kinematic features. When comparing data from a hand movement stopping task, there exist cortical sites specific for speech stopping and also sites involved in both speech and hand stopping, implying a range of inhibitory functions targeting unimodal and multimodal motor output (Breshears et al., 2019; Gavrilov and Nieder, 2021). Finally, we applied electrical stimulation and found that stimulation induced speech arrest overlapped with electrodes showing stop activity. These results provide novel evidence to support a previously unknown function of the premotor cortex in the inhibitory control of speech.

The inhibitory control of action has been studied extensively in the past decades. Results from neuroimaging studies, intracranial recordings, and stimulation experiments have supported a notion that the cognitive-level control of inhibition is largely confined to regions in the prefrontal cortex (Bari and Robbins, 2013; Rubia et al., 2003; Swann et al., 2009; Wessel et al., 2013). Specifically, the rIFC and pre-SMA are considered the core of the cortical control of response inhibition (Aron et al., 2004, 2014). Neural activity in the rIFC activates the subthalamic nucleus through a hyperdirect pathway to achieve so-called “outright” stopping in a typical stop-signal task (Aron and Poldrack, 2006; Aron et al., 2007; Chen et al., 2020). Our results on early stopping of speech provide novel findings that are complementary to the existing framework. First, although the spatial location of stop activity is partially overlapped with rIFC, it is primarily in the premotor regions, including many along the rostral part of the precentral gyrus (Fig. 1D). Second, the neural activation occurred bilaterally, with heavy involvement of the left hemisphere. While the difference in brain regions may be related to different task designs, e.g. canceling motor output at the initiation stage (Logan and Cowan, 1984) compared to stopping an ongoing behavior, the left hemisphere activity is likely to be vital for immediate interaction with the speech production network on the same side. Indeed, we found that the stop activity temporally preceded stop action, which has not been shown clearly in previous studies (Swann et al., 2013). The magnitude of stop activity also predicted the type of stopping required (i.e. midword or end-of-word), suggesting that specific articulatory patterns were affected during production. Third, previous studies found pre-SMA has an important role in inhibitory control (Chao et al., 2009; Chen et al., 2010; Swann et al., 2012). Here we observed pre-SMA activity, although weak in amplitude, in our speech task (Fig. 1D). However, our data suggests that the activity in the lateral premotor areas did not originate from pre-SMA (Fig. 5E, F). Our results and previous studies together suggest that there may be multiple sources for inhibitory signals in the cortex and stop activity in the premotor frontal cortex is one of them. Future work is needed to map the downstream pathway of premotor stop activity and probe whether aspects of stop activity are related to “proactive” stopping control described in previous studies (Chikazoe et al., 2009; Swann et al., 2013).

Our results also provide a novel view of speech arrest in clinical settings. The prevailing understanding is that speech arrest sites are indispensable nodes in the speech production network (Lu et al., 2021; Ojemann et al., 1989; Penfield and Roberts, 1959; Sanai et al., 2008; Tate et al., 2014). Stimulation disrupts the essential neural activity in these regions and therefore speech can no longer be generated. Using the same electrodes in the ECoG grids, we compared the location of speech arrest sites and the neural activity for production and stopping in our study. Surprisingly, we found that the spatial location of speech arrest sites did not overlap well with neural activity related to speech production or preparation, suggesting that disruption of motor related activity was not the sole cause of speech arrest. In contrast, we found a direct overlap between sites with speech arrest and those with stop activity. Therefore, stimulation may alternatively activate an inhibitory pathway of speech control, which contains the premotor regions where stop activity was found. This is supported by previous data in which the most common locations for speech arrest were found in the same premotor regions (Chang et al., 2017; Lu et al., 2021). The inhibition-based explanation of speech arrest has been speculated in the past (Ferpozzi et al., 2018; Mikuni et al., 2006; Rech et al., 2019; Silva et al., 2022). For example, negative motor areas (NMA) were reported across peri-rolandic and premotor regions, where stimulation induced cessation of movement (Filevich et al., 2012; Lu et al., 2021; Mikuni et al., 2006; Rech et al., 2019). It has been proposed that the function of NMAs may relate to inhibitory control rather than general motor programming (Filevich et al., 2012). Our results were consistent with this view. Another view suggests that speech arrest during right hemisphere stimulation may be caused by inhibitory control whereas speech arrest during left hemisphere stimulation may be due to disruption of speech programming or motor activity (Hannah and Aron, 2021). Here we found stop activity in both hemispheres, suggesting that inhibitory function may underlie speech arrest for both hemispheres. One interesting observation in our data was that certain areas in the precentral gyrus showed shared stop activity for both speech and hand movement. This is consistent with recent neurosurgical studies where stimulation in precentral sites induced both speech and hand arrest (Breshears et al., 2019; Leonard et al., 2019; Zhou et al., 2022). Localized regions with such effect may act on an executive control level where inhibition is generally considered multimodal (Cai et al., 2012; Wessel and Aron, 2017; Xue et al., 2008). However, speech-specific inhibition does exist within the ventral and middle premotor regions, albeit adjacent to sites with general inhibitory effects.

There are several reasons why speech arrest sites may not be critical for speech production. Speech arrest has been described bilaterally, and also involves the SMA regions. The right hemisphere and SMA lesions or resection do not lead to long-standing speech impairments. Furthermore, anarthria or pure speech arrest is not what one would expect from disruption of speech planning; instead paraphasias or apraxic speech would be expected. Re-defining speech arrests sites as general inhibitory rather than essential for production, has major clinical implications. For example, expanding surgical resection to include a speech arrest site may be tolerated and not results in speech production deficits.

Finally, our results have important implications for current models of speech production, particularly regarding the frontal cortex (Guenther, 2016; Hickok, 2012), notably the ventral part of the precentral gyrus (Bouchard et al., 2013; Chartier et al., 2018) and Broca’s area (Flinker et al., 2015; Hickok, 2012; Long et al., 2016). While these regions generally have faciliatory functions in speech production according to existing models, data from our study showed distinct but intermixed inhibitory functions in the same structures (see Fig. 1D). Despite this spatial overlap, some interesting differences exist regarding the electrode populations associated with these two functions. At the individual electrode level, electrodes with stop activity and those with speech production related activity did not largely overlap, indicating that the inhibitory and faciliatory functions may be separate at a smaller spatial scale than previously thought (Fig. 1E). This suggests that, instead of having separate anatomical regions for different functions, brain areas central to speech production may have mosaic subclusters of neural circuits. Furthermore, there is spatial segregation between electrodes showing stop activity and those encode AKT features (Fig. 5C, D). Given our initial evidence of directed functional connectivity (Fig. 5E) and the difference in the modulation of stop activity in midword trials (Fig. 4), it is highly likely that the inhibitory function of speech interacts with articulation and planning for speech production, perhaps in relation to the prosodic structure. Future studies should aim to delineate the interaction and further elucidate the connectivity within this speech inhibition network. Our results also imply that inhibitory control may play a more important role in speech production than previously thought. In verbal communication where there is a natural, fast exchange of phrases and sentences, inhibitory signals are likely to interlace with excitatory signals with precise timing. In speech impairments such as stuttering, aberrant inhibitory signals may cause the frequent stops found in production (Chang and Guenther, 2020; Eggers et al., 2010; Markett et al., 2016; Neef et al., 2018; Orpella et al., 2022). Investigations into the relationship of stop activity with normal and abnormal speech will be important future directions.

In summary, the results we describe suggest a previously unknown mechanism for the inhibitory control of speech production. We provided a framework for a deeper understanding of the neural process underlying speech production with important implications for both basic neuroscience and clinical practice.

## Acknowledgments

We thank Patrick Hullett and others for their help with electrocortical stimulation. We thank John Rolston for the initial work on the project. We thank Matthew Leonard and Sarah Harper for their comments on the manuscript. We thank Richard Ivry, Junfeng Lu, Xiuli Tong for helpful discussions on the results. We thank Jonathan Kleen, Neal Fox, Maansi Desai and Alia Shafi for help with the intra-operative mapping data analysis. We thank Anthony Fong for technical assistance and record inquiries. We thank Yuanning Li for suggestions on statistics. This work was supported by grants from the NIH (DP2OD008627 and U01NS098971-01 to E.F.C, and K99DC020235 to L.Z.). This research was also supported by the New York Stem Cell Foundation, the Howard Hughes Medical Institute, the McKnight Foundation, the Shurl and Kay Curci Foundation, and the William K. Bowes Foundation.

## Author Contributions

L.Z. and E.F.C. conceived the research and designed the experiment. L.Z., E.F.C., and others collected the data. L.Z., A.B.S., and G.L.K. analyzed the data. L.Z., A.B.S., and E.F.C. wrote and revised the manuscript. E.F.C. supervised the project.

## Methods

### Participants

Eleven participants (3 male, 8 female; mean ± std. age: 31 ± 9 years) performed a speech stopping task and were included in the analysis. Five additional participants (1 male, 4 female; mean ± std. age: 35 ± 13 years) were included in the analysis for the sentence reading task. All participants were patients undergoing intracranial monitoring for intractable epilepsy. Of the participants in the speech stopping task, 8 had left hemisphere coverage and 3 had right hemisphere coverage. For the additional participants involved in the sentence reading task, 3 had left hemisphere coverage and 2 had right hemisphere coverage. All procedures and protocols were approved by the Institutional Review Board of the University of California, San Francisco. Participants provided written informed consent before participating in the studies. All participants were fluent English speakers and had no cognitive deficits that could potentially affect the study.

### Task design and setup

To test early stopping, participants performed a speech stopping task where they followed visual cues to start and stop speaking (Fig. 1A). The task was custom-programmed using Psychtoolbox-3 and MATLAB to be presented on a laptop screen (Microsoft Surface Book 2). Each trial started when a green circle (go cue) was presented at the center of the screen and participants were instructed to start reciting the days of the week. After a random delay (2-5 s), the circle changed to red color (stop cue) and participants were instructed to stop immediately. The time between the stop cue and the acoustic stop of speech is referred to as “stop reaction time” (SRT). The next trial started after a short pause (2.5-3.5 s), and participants continued with recitation when the circle turned green. Throughout the trial, a cross was shown at the center of the screen inside the colored circle, and the participants were asked to fixate on the cross. To synchronize with neural recordings, a gray rectangle flashed at the upper right corner of the screen at the same time as the presentation of cues. A photodiode (S2281-01, Hammamatsu) was placed at the location of the rectangle on the screen and was connected to the neural recording setup. A small number of participants recited the months of the year or counted continuously based on preference.

To compare early stopping against a natural finish condition, a subset of participants (N = 4 out of 11) performed a sentence reading task (Chartier et al., 2018). Briefly, participants read aloud 100 sentences from the MOCHA-TIMIT database (Wrench, 1999). One sentence was presented on the screen in each trial and participants read the sentence at their own pace. There were no cues to indicate when they should stop. Each sentence was read once so a total of 100 trials were performed. Additional data with the same sentence reading task collected from previous studies (N = 5 participants, 100 trials for each participant) were also included.

To compare the neural activity evoked by stopping speech to stopping hand movements, a subset of participants (N = 8 out of 11) performed a hand movement stopping task. This task was modified from the speech stopping task with a similar presentation of cues. When the go cue was presented, participants were instructed to push a button repetitively and rhythmically using their contralateral thumb in reference to the ECoG grid. At the presentation of the stop cue, they were instructed to stop and release the button immediately. The time delay between the go cue and the stop cue was randomly jittered between 2-4 s.

### Data acquisition and preprocessing

Electrocorticography (ECoG) was acquired through subdural high-density grids (Integra or AdTech) with 1.17 mm diameter exposed contacts and 4 mm center-to-center spacing. The voltage time series signal (raw signal) from each electrode contact was amplified and digitized at a sampling rate of 3051.7578125 Hz by a pre-amplifier (PZ5, Tucker-Davis Technologies) and processed through a digital signal processor (RZ2, Tucker-Davis Technologies). To obtain the high-gamma analytic amplitude (HGA), raw signals were down-sampled to 400 Hz, notch-filtered at 60, 120, and 180Hz to remove line noise, and Hilbert-transformed at 8 logarithmically distributed bands within 70-150 Hz. Each of the 8 bands was z-scored and the average was taken between bands to obtain one HGA time series. Raw signals and HGA were visually inspected to exclude bad channels and trials with artifacts. Speech audio from participants was recorded through a microphone (e845-S, Sennheiser), amplified by a microphone amplifier (MA3, Tucker-Davis Technologies), and digitized through a digital signal processor (RZ2, Tucker-Davis Technologies). The output of the push-button device (voltage signal) was digitized and recorded (RZ2, Tucker-Davis Technologies).

### Electrode localization

A pre-operative MRI and a post-operative CT scan were used to register the location of electrodes relative to the brain. Reconstruction of the pial surface was performed using the pre-operative MRI image in Freesurfer. To convert the electrode coordinates from individual brains to the average brain (cvs_avg35_inMNI152 template), we used a nonlinear surface registration in Freesurfer with a spherical sulcal-based alignment (Fischl et al., 1999). This method ensures that electrodes on a given gyrus in the original participant’s space remain on the same gyrus in the converted space. It does not, however, maintain the original geometry of the electrode grid.

### Data analysis on stop activity and production activity

We identified the acoustic start and stop time of speech production in each trial using an energy threshold-based method. If necessary, further adjustment was made by visualizing the spectrogram of the microphone signals. To avoid the condition where participants stopped speaking by themselves independent of the stop cue, we excluded trials with SRT less than 0.1 s (Fig. 1B). To test if an electrode shows stop activity, we used a baseline period [-0.5,0]s before the stop cue and an analysis period [0,1]s after the stop cue. We set the baseline period to be during speech production because we intended to identify additional activation beyond the neural modulation caused by production. Since activity may show different delays after the stop cue across electrodes, we used a series of overlapping sliding windows of 0.2s duration (0.1s step size) within the analysis period. We compared the mean HGA in each of these windows with respect to the mean HGA in the baseline period and tested significance with a paired t-test. Significant activation is determined if the mean HGA in the sliding window is larger than that in the baseline period and if the p-value is smaller than 0.0001. To control for multiple comparisons, we used an FDR correction for each sliding window across electrodes within each participant at a q-level of 0.05. An electrode is regarded as showing significant stop activity if any of the sliding windows showed significant activation. To test activity during speech production, we used a baseline period of [-0.5, 0]s before the go cue and an analysis period of [-1.5, -0.5]s before the stop cue. This analysis period covered the late part of speech production in most trials. We chose these analysis intervals to exclude electrodes that were only activated during the initial part of speech production. Applying this method allowed us to select electrodes that showed sustained activation throughout production. To compare HGA in the analysis period with that in the baseline period, we used a similar series of sliding windows as for stop activity and we followed the same steps for statistical tests. To summarize the magnitude of activity (Fig. 1D, E), we used the maximum t-value among the significant sliding windows.

### Comparison with the natural finish condition

For each individual electrode, we compared the activity at the end of speech production in the sentence reading task with the activity in the speech stopping task (Fig. 2A, B, D, E). Since there was no stop cue in the sentence reading task, we used a [-0.5, 0.5] s analysis period centered at the time of speech stop. Again, we applied sliding windows of 0.2s (0.1s step size) duration within the analysis period. We used a baseline period [-1, -0.5]s relative to the speech stop. A similar strategy to identifying the stop activity in the speech stopping task is adopted here to test for significant activation during a natural finish. The magnitude of activation is quantified by t-value, which is taken from the max t-value among the significant sliding windows. To delineate whether the activity at the end of speech production is associated with regular motor activity, we divided the electrodes into two groups according to whether they have production activity in the speech stopping task, as shown in Fig. 1. To compare the overall activity between early stopping and natural finish conditions, we included data from 5 additional participants that performed the same sentence reading task for previous research. These additional participants had similar electrode coverage as those in the current study.

### Temporal correlation of stop activity

To test whether stop activity in a given electrode was more correlated with the stop cue or stop action, we performed single-trial timing analyses. Because HGA in single trials is noisy, taking the maximum does not always capture the correct timing when the majority of HGA activation occurred. In the following analysis, we calculated the “dominant activity time” and used it as an alternative measure of the peak time.

To identify stop action related electrodes, we first aligned each trial to speech stop and calculated the average HGA. We performed a cross-correlation between the average HGA and the single-trial HGA and identified the peak of the cross-correlation. We then used a threshold to include trials with reasonably large peaks in the cross-correlation. This threshold was set at the 95th percentile of the amplitude of all cross-correlation results. Only trials with cross-correlation peaks higher than this threshold were included and were assigned with the dominant activity time. Dominant activity time was calculated as the peak of the average HGA plus the offset obtained by cross-correlation. We then fit linear and quadratic models between the dominant activity time and the stop reaction time (the linear model is equivalent to Pearson correlation). If either model fit showed significance (p < 0.05) and if the correlation coefficient was positive for the linear model, then this electrode was determined as a stop action related electrode.

To identify cue related electrodes, we first aligned the trials to the stop cue and obtained the averaged HGA. We then calculated the dominant activity time using cross-correlation, similar to the previously explained technique for stop action related electrodes. We measured the difference between the dominant activity time and stop reaction time and then calculated the Pearson correlation between this difference and the stop reaction time. If the correlation is significant (p < 0.05), the standard deviation of the dominant activity time is smaller than 0.15s, and the electrode is not stop action related, then it is determined as a cue related electrode. Electrodes not meeting the criteria for either type of alignment were assigned a label of “other”.

To quantify the activation start time relative to the time of speech stop (Fig. 3F), we aligned the HGA to the speech stop and used sliding windows (0.2s duration, 0.1s step size) within [-0.5, 0.5]s period centered at speech stop. Similar to the previously described methodology, we tested whether each of these windows had a significant increase in HGA compared to the baseline period ([-0.5,0]s relative to the stop cue). We used the center of the window that had the earliest significant activation (p < 0.05 for paired t-test) as the activation start time. Multiple comparisons is controlled by an FDR correction for each sliding window across electrodes within each participant at a q-level of 0.05.

### Low-frequency activity

Beta-band activity was used in the comparison of early stopping and natural finish conditions (11 participants for early stopping; 9 participants for natural finish). To extract beta-band activity, the raw signal from each electrode was first notch filtered to remove line noise and then zero-phased filtered (using the “filtfilt” function in MATLAB) between 20-30 Hz. The analytic amplitude of the signal was next computed by taking the Hilbert transform on the filtered data. The analytic amplitude was then z-scored using periods of silence as the baseline. This analytic amplitude was used for the remainder of the analyses. To test if an electrode had a significant increase in beta-band activity during stopping, we compared the mean amplitude in an analysis window [0, 0.5]s relative to the time of speech stop to that in a baseline window [-0.5, 0]s relative to the stop cue. Such intervals were chosen based on previous studies on beta suppression during motor actions (Cassim et al., 2001; Sanes and Donoghue, 1993). Significance was determined using a paired t-test with FDR correction at a q-level of 0.05.

Delta-band activity (1-4 Hz) was extracted from the raw signals, similar to the procedure for the beta-band activity. To find significant activation in the delta-band activity, we compared the mean analytic amplitude in a 0.5s window directly after the stop cue to that in a 0.5s window directly before the stop cue. We used the same statistical criteria to determine the significance as for the beta-band activation.

### Comparison between midword and end-of-word trials

We labeled the production in each trial for whether the ending word was completed. If speech stopped before finishing the entire word, the trial was labeled as a midword trial. Otherwise, it was labeled as an end-of-word trial. We only included electrodes with stop activity but no production activity to avoid potential confound from signals reflecting any specific modulation to articulators during stopping. We compared the average HGA aligned to the speech stop between the two types of trials using a cluster-based permutation rank-sum test. We used N = 1000 permutations and a total of one cluster as the parameters. The cluster-based permutation test corrected for multiple comparisons when performing timepoint-by-timepoint comparisons between the two types of trials. To visualize the size of the stop type effect (Fig. 4C), we took the largest z-value of the rank-sum test within the significant cluster of each electrode. To characterize when the activity started to show the difference between the two stop types, we used the earliest time point in the significant cluster.

For the population analysis, we built a linear classifier for each participant to predict whether a single trial is a midword or end-of-word trial. Specifically, we used the average HGA within a 0.2s window centered at speech stop from all stop electrodes without production activity. We used an L1-regularized logistic regression for the classifier. We performed 500 repeats, each time randomly picking 70% data as the training set (30% as the testing set), to obtain the mean and 95% confidence interval of the performance. We used the area under the ROC curve (AUC) to quantify the classifier performance.

### Encoding of articulatory kinematic trajectory (AKT)

We followed similar steps as described previously (Chartier et al., 2018) to fit AKT encoding model for the frontal and parietal electrodes using 13 total features. We used an acoustic-to-articulatory inversion (AAI) algorithm (Parrot et al., 2020) to obtain the X and Y coordinates of 6 vocal tract points during speaking. These vocal tract points included the tongue dorsum, tongue blade, tongue tip, lower incisor, upper lip, and lower lip. We also included the fundamental frequency (F0) which represented the laryngeal feature. F0 was calculated using the “pitch” function in MATLAB on voiced phonemes. In the other part of the speech, F0 was set to zero. HGA of each electrode was fit with a linear encoding model, which predicted the HGA as the convolution of articulator kinematics (13 dimensions) with a temporal filter (Fig. 5B). We chose a filter width of 0.5s, given results from the previous study (Chartier et al., 2018). The model was fit using ridge regression on a training set composed of 80% of the data. The ridge parameter was evaluated with a 20-way bootstrap procedure based on the training set for each electrode. The final ridge parameter was chosen as the optimal value using the average across all electrodes. Pearson’s correlation coefficient, r, calculated between the model predicted activity and the actual HGA, was used to evaluate the model fit based on a testing set composed of 20% of the remaining data. Electrodes with a strong encoding of AKT features are determined as those with r > 0.2.

### Granger causality

To quantify the functional connectivity between brain regions, we calculated the Granger causality (GC) for pairs of electrodes using the MVGC toolbox (Barnett and Seth, 2014). Raw signals were first notch filtered to remove line noise and then used in the GC analysis. We used the autoregressive integrated moving average (ARIMA) model to pre-whiten the signal to meet the stationarity requirement (Leuthold et al., 2005). A Kwiatkowski-Phillips-Schmidt-Shin (KPSS) test was then applied to confirm the signal was stationary. We computed GC in the frequency domain between each electrode pair from target regions. We took the max GC across the frequency spectrum as the GC between the electrode pair. To test whether an electrode pair has significant functional connectivity, we shuffled data across trials and calculated GC on the shuffled data to obtain a “null distribution” (N = 500 permutations). Electrode pairs with GC values within or larger than the top 0.2% of the null distribution were considered significant and included in the analysis. We repeated the calculation in three time windows: a baseline window [-0.5, 0]s relative to stop cue; window 1 [0,0.5]s, and window 2 [0.5,1]s relative to stop cue. To compare whether there is a change in GC across electrode pairs between brain regions, we plotted the mean and standard error of all significant electrode pairs across the three windows (Fig. 5E). We used a repeated measures ANOVA with post-hoc analysis to find a significant change in GC between the windows (p < 0.05).

### Stopping of hand movement

We excluded all trials where hand stopping occurred before the stop cue. We followed the same sliding window based strategy as in the speech stopping task to identify the stop activity and the activity during hand movement. For each electrode, we compared the stop activity between the speech and hand movement tasks. We defined a selectivity index (SI). SI = (*t*_speech_ - *t*_hand_) / (*t*_speech_ + *t*_hand_), where *t*_speech_ is the t-value of the stop activity for speech and *t*_hand_ is the t-value of the stop activity for hand movement. If there was no significant stop activity for either modality, the corresponding t-value was set to zero. This index normalized the difference of activity to a range of [-1,1]. An SI of +1 indicated that the electrode only showed stop activity in speech and an SI of -1 indicated that the electrode only showed stop activity in hand movement.

### Electrocortical stimulation and speech arrest

Participants underwent electrocortical stimulation (ECS) as part of the clinical procedures to map functional areas critical for sensorimotor and language processing. This procedure provided an opportunity to compare the recorded neural activity (e.g., stop activity) during tasks with stimulation effects using the same electrodes. Bipolar current stimulation was delivered through a pair of electrodes by a clinical stimulator (Nicolet Cortical Stimulator, Natus Medical Incorporated, Fig. 7B). We used stimulation current characterized by a biphasic pulse train, typically with 50Hz frequency, 2s duration, and 2-6mA amplitude. Participants were instructed to count continuously or recite the days of the week, and stimulation was delivered during this ongoing process. The amplitude of the current was first set to be 2mA and gradually increased to probe whether stimulation affected speech production, induced sensation, or generated motor output. Speech arrest sites were tested multiple times to confirm findings.

The calculation of stop activity and production activity is the same as previously described. To test whether stimulation disrupted activity involved in planning, we calculated the pre-speech activity in the speech stopping task. Activity in an analysis period prior to speech production (between go cue and production onset) was compared to the activity in a baseline window (prior to go cue). Significant activation is tested with a paired t-test with FDR correction at a q-level of 0.05.

For an independent cohort of patients, we performed ECS during intra-operative mapping. These patients underwent ECS to identify essential sensory, motor, and language sites located in the lateral portion of the left hemisphere. ECS took place after craniotomy during which patients were gradually awakened by reducing their sedation. Stimulation was performed using a clinical stimulator (Ojemann Cortical Stimulator, Integra LifeSciences) with typical settings (60 Hz, bipolar, biphasic, 1ms pulse width). The stimulation threshold (range= 1.5-4.5 mA) was titrated on a per-patient basis to achieve the maximum possible stimulation current without causing after-discharges as determined by intraoperative electrocorticography. Patients were instructed to count slowly from one to thirty or recite the days of the week or months of the year. Stimulation was administered by the surgeon during the counting or recitation process. When stimulation was found to disrupt speech at a given cortical site, the site was stimulated nonconsecutively at least two more times, although the error replication protocol varied for a minority of patients. During ECS, the surgeon typically demarcated the locations of sensory, motor, and language sites with sterile paper tags. ECS mapping was video recorded (including capturing both the exposed brain and, with a second camera, the patient’s face) for later neuroanatomical co-registration and behavioral analysis. During data analysis, these videos were annotated to identify precise neuroanatomical loci on a given patient’s brain for each stimulation site. The stereotactic coordinates and gross anatomical region of interest for each stimulation site were recorded and registered as coordinates on a standardized, average brain using a custom MATLAB script by a trained neurologist blinded to the stimulation effect observed at each site.

## Supplementary Text

In addition to high-gamma band and beta-band activity, we also characterized delta-band activity during early stopping, which has been found to be important in response inhibition in previous studies (Chen et al., 2020; Huster et al., 2013). Figure. S2B shows an example electrode for which the averaged delta-band analytical amplitude increased after the stop cue when trials were time aligned to the stop cue. Similar increases in delta-band activity were found in other electrodes across participants (Fig. S2D). In our study, the electrodes with delta-band activation were mostly located in the middle and dorsal premotor cortex on the lateral side (Fig. S2D). On the medial side, the location of these electrodes partially overlapped with pre-SMA and was anterior to where we observed high-gamma activation.

---

**Fig. S1.**
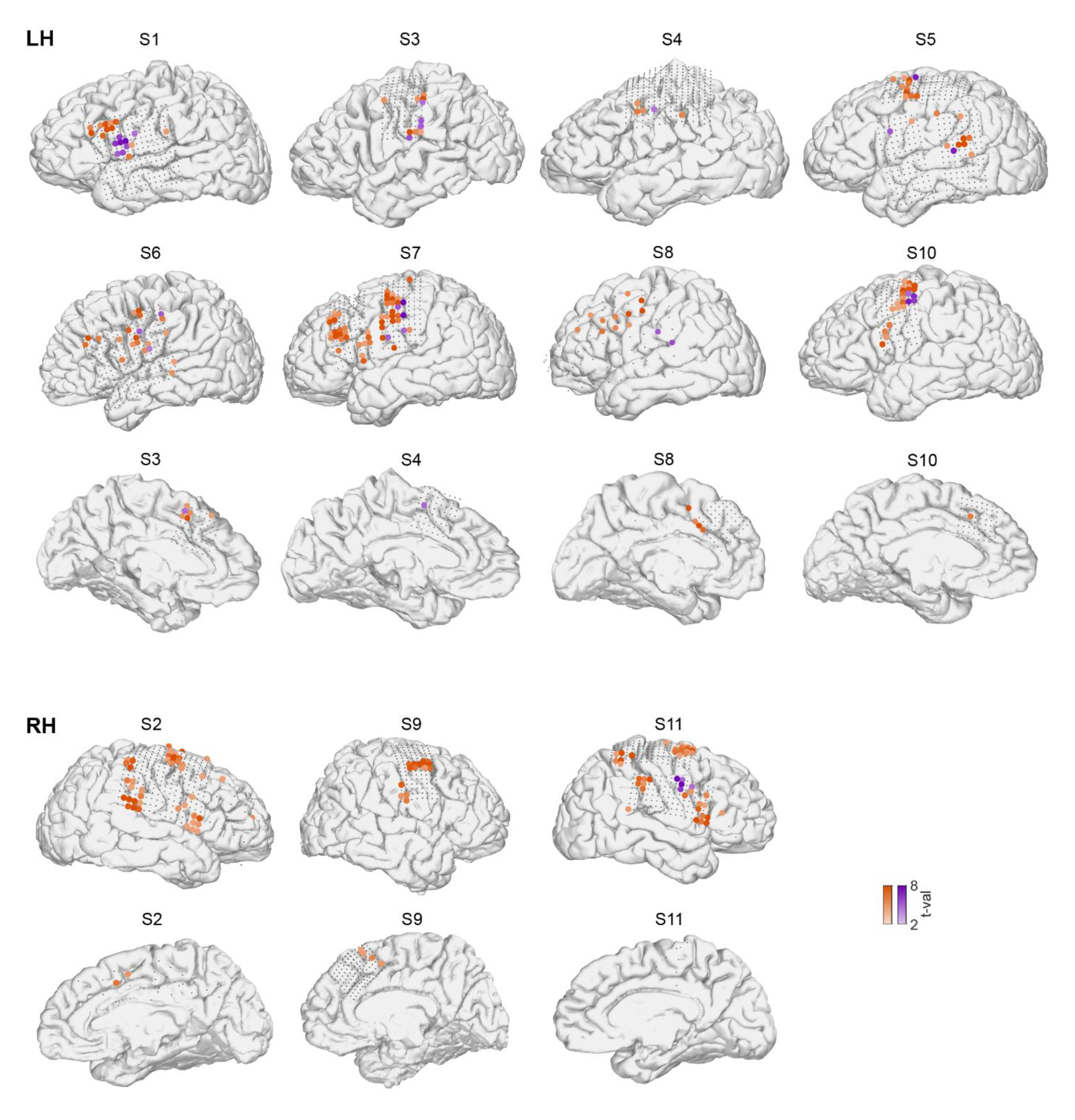
Stop activity for each participant. The spatial location of electrodes showing stop activities (colored circles) in each participant. Color intensity indicates the magnitude of activity (t-value). Orange: electrodes with no production activity. Purple: electrodes with production activity. Gray dots indicate the electrode coverage. LH: left hemisphere. RH: right hemisphere.

**Fig. S2.**
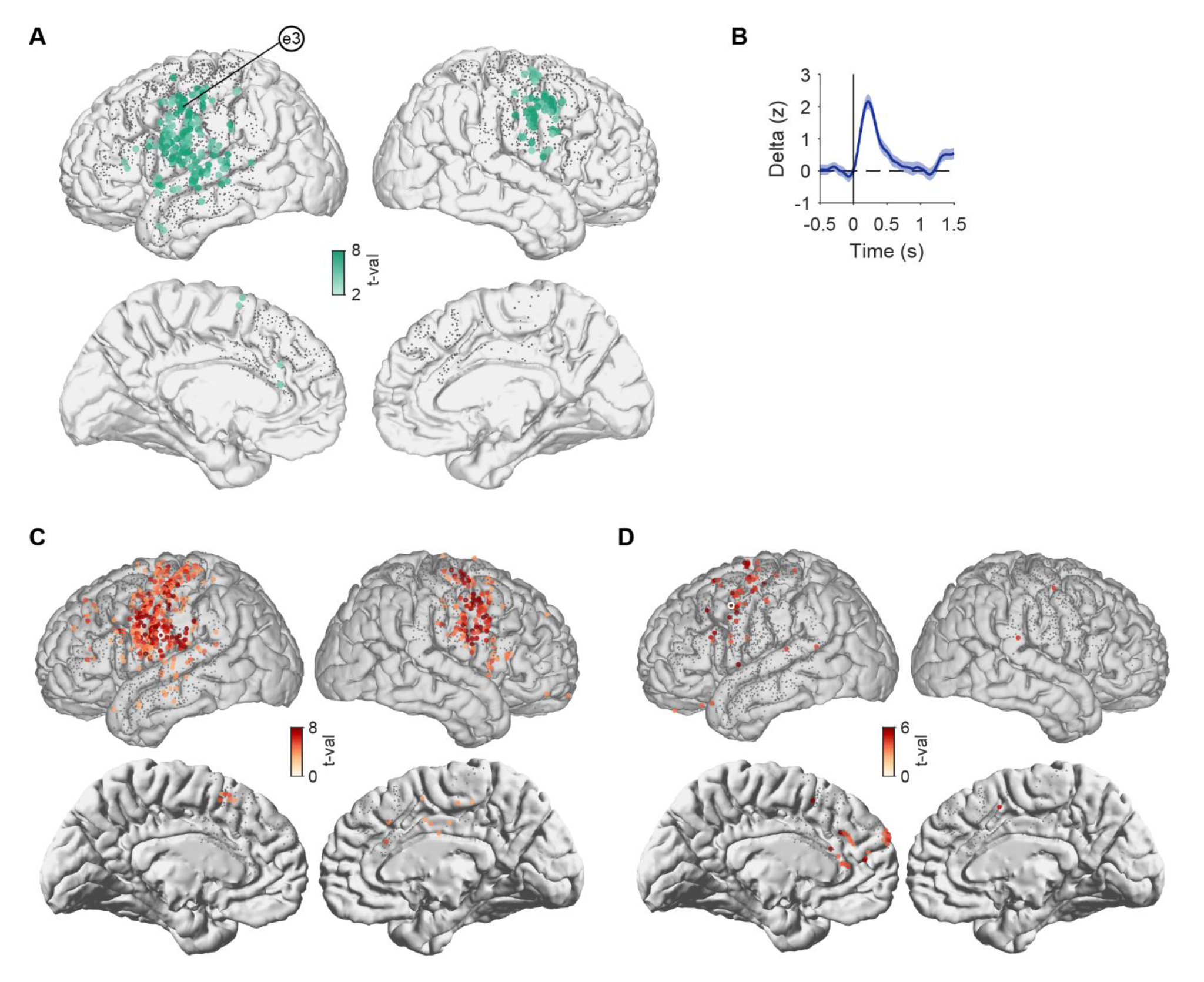
High-gamma activity during speech production and low-frequency activity during speech stopping. (A) Location of electrodes showing production activity from all participants plotted on an average brain. Color intensity indicates the magnitude of activity (t-value). (B) Averaged delta-band analytical amplitude (mean ± sem) in an example electrode, time-aligned to the stop cue. Time zero is the stop cue. (C) The spatial location of electrodes showing beta-band activation during stopping for all participants. Color intensity indicates the magnitude of activity (t-value). Gray dots indicate the electrode coverage. The white circle indicates the example electrodes shown in Fig. 2G. (D) Similar format to (C), for electrodes with delta-band activation. The white circle indicates the example electrodes shown in (B).

**Fig. S3.**
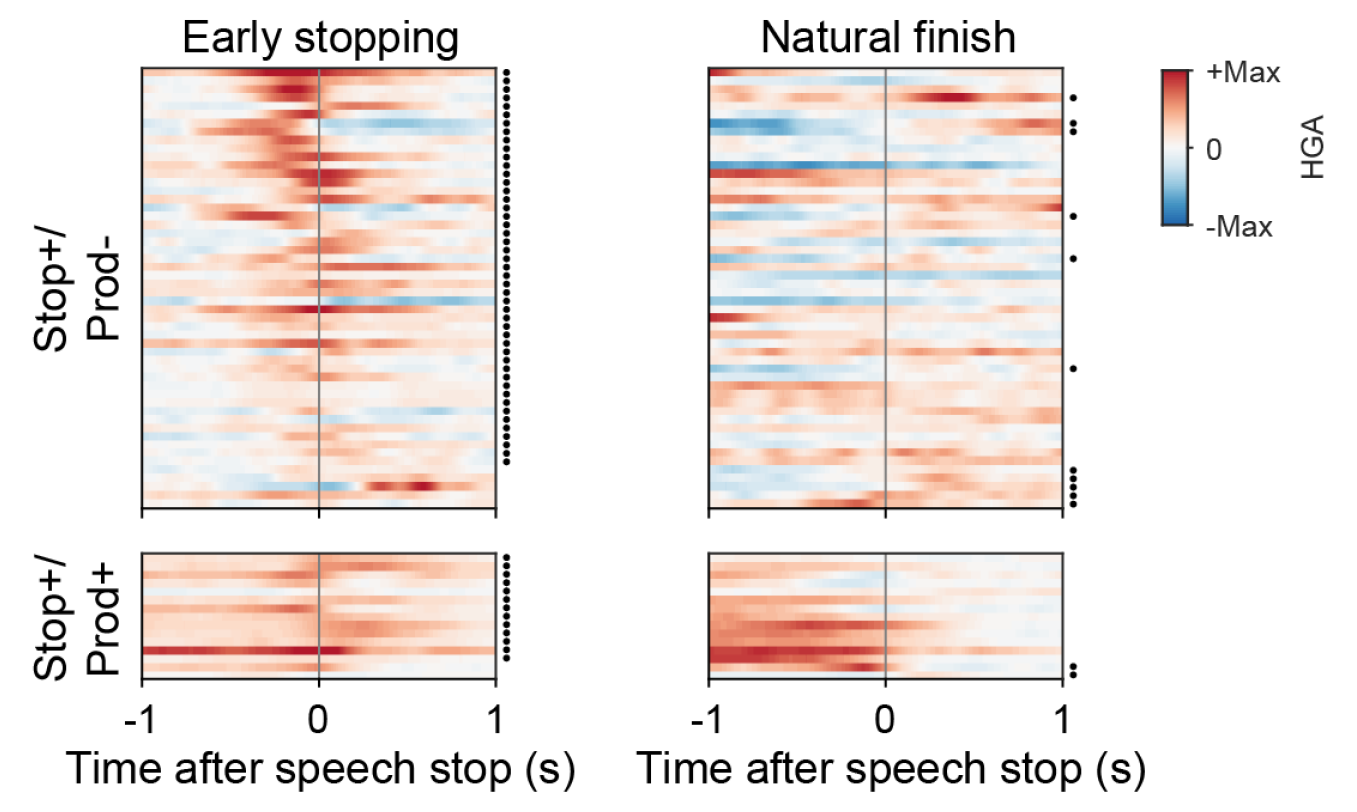
Comparison of high-gamma activity for individual electrodes between early stopping and natural finish conditions. The heatmaps illustrate the averaged activity across trials for each electrode. Each row indicates one individual electrode, with the same electrode order in the left and right panels. Black dots to the right of the heatmap indicate significant stop activity.

**Fig. S4.**
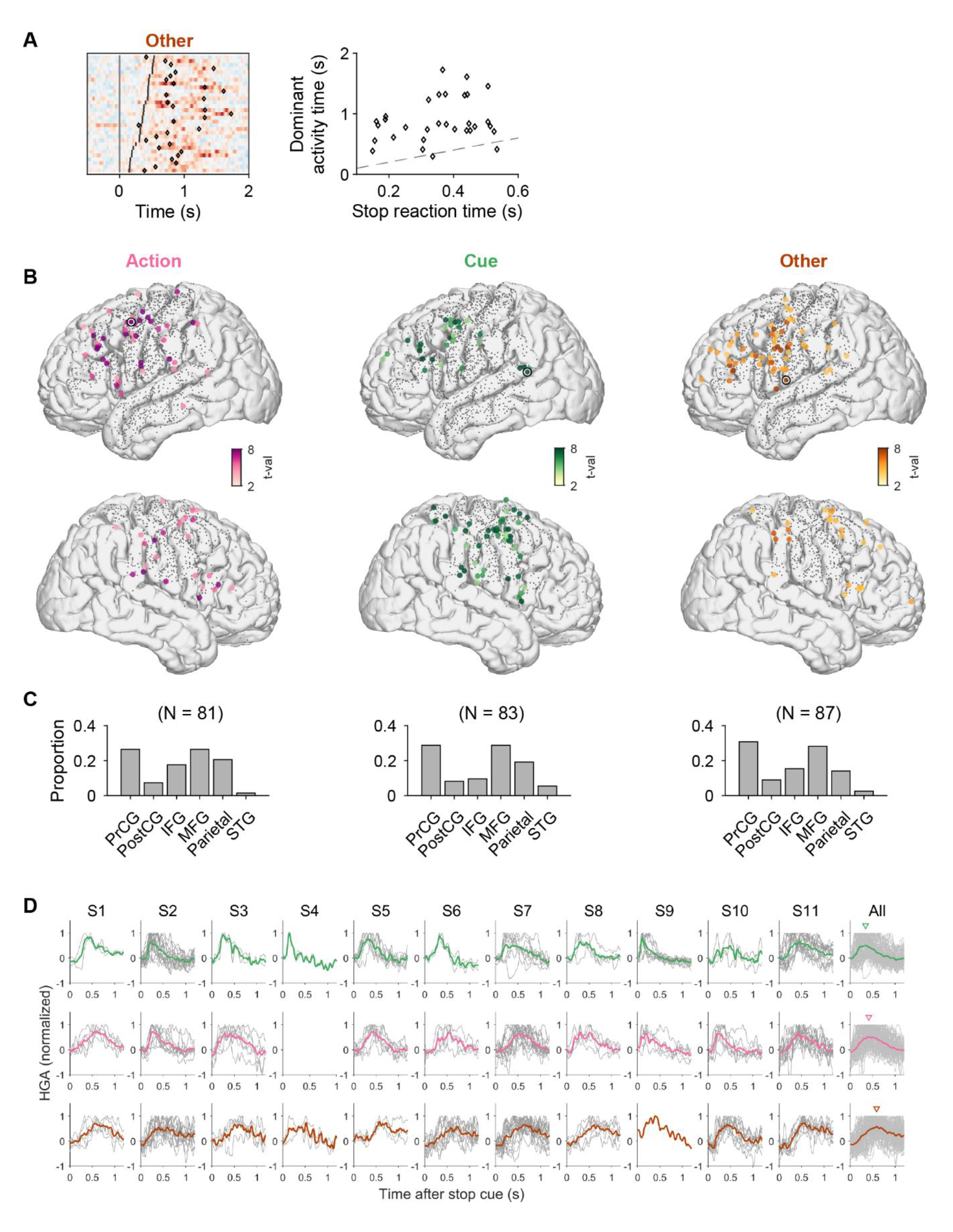
Spatial location of electrodes showing action related, cue related, and other activity. (A) Similar to Fig. 3A, C, example electrode showing stop activity not correlated to either stop action or stop cue (referred to as “other”). (B) Location of electrodes belonging to the three types. The black-and-white circles indicate the location of example electrodes shown in Fig. 3A-D and in (A). (C) The proportion of electrodes found in each brain region. (D) Stop activity from each participant grouped by the types. Gray curve: normalized activity from individual electrodes. Thick colored curve: averaged activity from all electrodes within individual participants (S1-S11) and for all participants (rightmost column). Downward triangles indicate the peak time of the smoothed data shown in the colored curve.

**Fig. S5.**
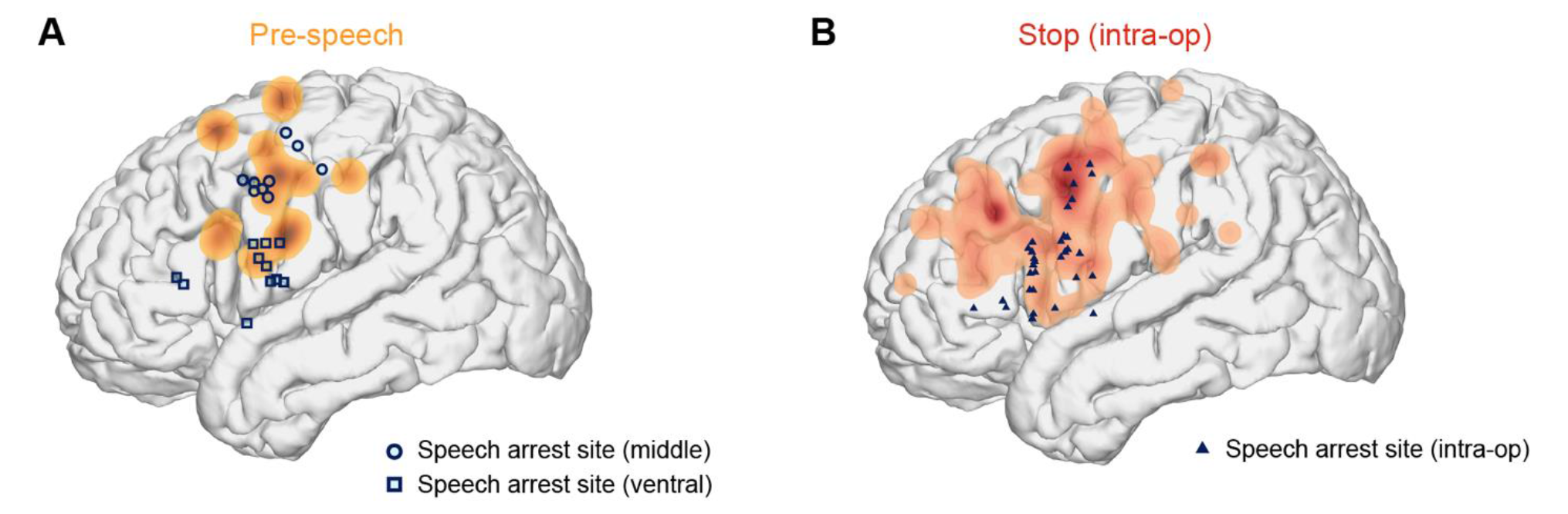
Comparison of the spatial location of speech arrest sites and neural activity. (A) Density map of electrodes with pre-speech activity but no stop or production activity, overlaid with speech arrest sites. (B) Similar to Fig. 7C left panel, the same density map of stop activity, overlaid with speech arrest sites found in a separate dataset of intra-operative mapping.

